# Sensitive cluster-free differential expression testing

**DOI:** 10.1101/2023.03.08.531744

**Authors:** Alsu Missarova, Emma Dann, Leah Rosen, Rahul Satija, John Marioni

## Abstract

Comparing molecular features, including the identification of genes with differential expression (DE) between conditions, is a powerful approach for characterising disease-specific phenotypes. When testing for DE in single-cell RNA sequencing data, current pipelines first assign cells into discrete clusters (or cell types), followed by testing for differences within each cluster. Consequently, the sensitivity and specificity of DE testing are limited and ultimately dictated by the granularity of the cell type annotation, with discrete clustering being especially suboptimal for continuous trajectories. To overcome these limitations, we present miloDE - a cluster-free framework for differential expression testing. We build on the Milo approach, introduced for differential cell abundance testing, which leverages the graph representation of single-cell data to assign relatively homogenous, ‘neighbouring’ cells into overlapping neighbourhoods. We address key differences between differential abundance and expression testing at the level of neighbourhood assignment, statistical testing, and multiple testing correction. To illustrate the performance of miloDE we use both simulations and real data, in the latter case identifying a transient haemogenic endothelia-like state in chimeric mouse embryos lacking Tal1 as well as uncovering distinct transcriptional programs that characterise changes in macrophages in patients with Idiopathic Pulmonary Fibrosis. miloDE is available as an open-source R package at https://github.com/MarioniLab/miloDE.

## Introduction

Diseases are complex and dynamic processes, with different organs, tissues, and cell types undergoing unique changes that often manifest at the transcriptional level. Single-cell genomics provides a sensitive and unbiased lens for identifying how cell type specific phenotypes change following a perturbation or in disease (Tirosh et al. 2016; Stephenson et al. 2021; Rood et al. 2022). To perform a comparative analysis between conditions, it is necessary to employ a case-control experimental design that requires the existence of several replicates from the control (typically healthy or wild type samples) condition together with replicates with the case phenotype (typically disease or perturbation). The standard workflow to identify potential case-specific phenotypes involves the batch-corrected embedding of all samples on the same low-dimensional space followed by comparative analysis between ‘comparable’, transcriptionally similar cells (e.g. within the same cell type) (Hao et al. 2021; Lotfollahi et al. 2022; Dann, Teichmann, and Marioni 2022). Examples of such comparative analyses include differential abundance (DA) testing - a framework to estimate whether different cell types change their relative abundance between conditions, and differential expression (DE) analysis, in which individual genes that are expressed at different levels between conditions are identified.

DE analysis, at both the bulk and single-cell level, has yielded valuable insights into the mechanisms of numerous diseases (X.-S. Wang et al. 2006; Adams et al. 2020; Elmentaite et al. 2020) and enabled the identification of drug targets (Montoro et al. 2018; van den Hurk et al. 2022). Accordingly, numerous methods for differential expression testing have been proposed and benchmarked (Soneson and Robinson 2018; T. Wang et al. 2019; Crowell et al. 2020; Squair et al. 2021; Gagnon et al. 2022). Methods originally developed for bulk RNA-seq, such as limma-voom (Ritchie et al. 2015), edgeR (Robinson, McCarthy, and Smyth 2010; McCarthy, Chen, and Smyth 2012; Y. Chen, Lun, and Smyth 2016), and DESeq2 (Love, Huber, and Anders 2014), were adapted for scRNA-seq analysis, by aggregating mRNA counts from transcriptionally similar cells, yielding so-called “pseudo-bulk” counts. As scRNA-seq has become increasingly popular, methods designed to handle specific properties of scRNA-seq data have been proposed (SCDE, scDD, D3E, MAST, DECENT, etc) (Kharchenko, Silberstein, and Scadden 2014; Korthauer et al. 2016; Delmans and Hemberg 2016; Finak et al. 2015; Ye, Speed, and Salim 2019). The main factor distinguishing these methods is the assumed underlying distribution of the expression counts (the main being the Poisson and the Negative Binomial, as well as the zero-inflated versions of those). Finally, a recently proposed approach, lvm-DE, uses a Bayesian framework leveraging posterior distributions estimated from deep generative models, which is suggested to be suitable for the complex, non-linear experimental designs that are particularly relevant for performing DE analysis between groups in extensive cohort studies with complex metadata (Boyeau et al., n.d.).

Importantly, all of these methods require that cells are grouped into presumably homogenous, transcriptionally similar clusters (i.e. cell types). This minimises variability in gene expression counts between cells within a sample, mitigates the inflation of p-values and increases the power to detect differential expression for lowly expressed genes (Lun and Marioni 2017). However, performing statistical tests at the cell type level (i.e. asking whether a gene is differentially expressed between conditions within a specific cell type) is ultimately limited and dictated by the sensitivity and resolution of the cell type annotation, which is a highly subjective and study-dependent process. As a result, genes that are DE only within a certain homogenous sub-region within the cell type (with the cell type either consisting of several discrete subpopulations, continuous trajectories, or a combination of both), will often be undetected. On the other hand, if sub-cell type composition differs between case and control samples, genes that are specific in their expression to a local sub-region of a cell type (in both conditions) might get falsely identified as showing significant DE when, in fact, they ‘mark’ DA sub-regions. Finally, due to potentially substantial differences in cell type abundances, the power to identify statistically significant differences in expression can vary across cell types.

These shortcomings motivate the development of frameworks that are more sensitive to the local transcriptional structure, and that will learn the per gene DE on the manifold rather than separately for each annotated cell type. For example, the recently proposed computational suite Cacoa incorporates a DE framework that is performed at a single-cell level (Petukhov et al. 2022). However, it does not account for potential confounding covariates and batch effects, making it challenging to apply in many settings, including cohort studies and complex experimental designs which are becoming increasingly popular.

To overcome these limitations, we present miloDE - a cluster-free framework for DE testing. We extend Milo, a method for cluster-free differential abundance (DA) testing, where cell abundance is estimated over overlapping neighbourhoods on the kNN graph representation of scRNA-seq data (Dann et al. 2021). We address key differences between DA and DE testing at both the neighbourhood assignment level and when performing the statistical testing and multiple hypothesis testing correction, allowing us to perform DE detection for each gene and each neighbourhood. Importantly, the fine neighbourhood resolution of miloDE unlocks a suite of methods tailored for scRNA-seq analysis, enabling the detection of co-regulated transcriptional modules containing genes that change their expression in a coordinated manner. Finally, we demonstrate the performance of miloDE in both simulated data and in different biological contexts. miloDE is an open-source R package, available at https://github.com/MarioniLab/miloDE.

## Results

### Overview of the method

miloDE is a cluster-free framework for DE testing that leverages overlapping neighbourhoods in the graph representation of scRNA-seq data (Fig. 1). Once the graph is constructed, we select a subset of cells (referred to as index cells); subsequently, each index cell along with its neighbours is assigned to a single neighbourhood (see Methods). Once neighbourhoods are assigned, DE testing is carried out within each neighbourhood.

**Figure 1.**
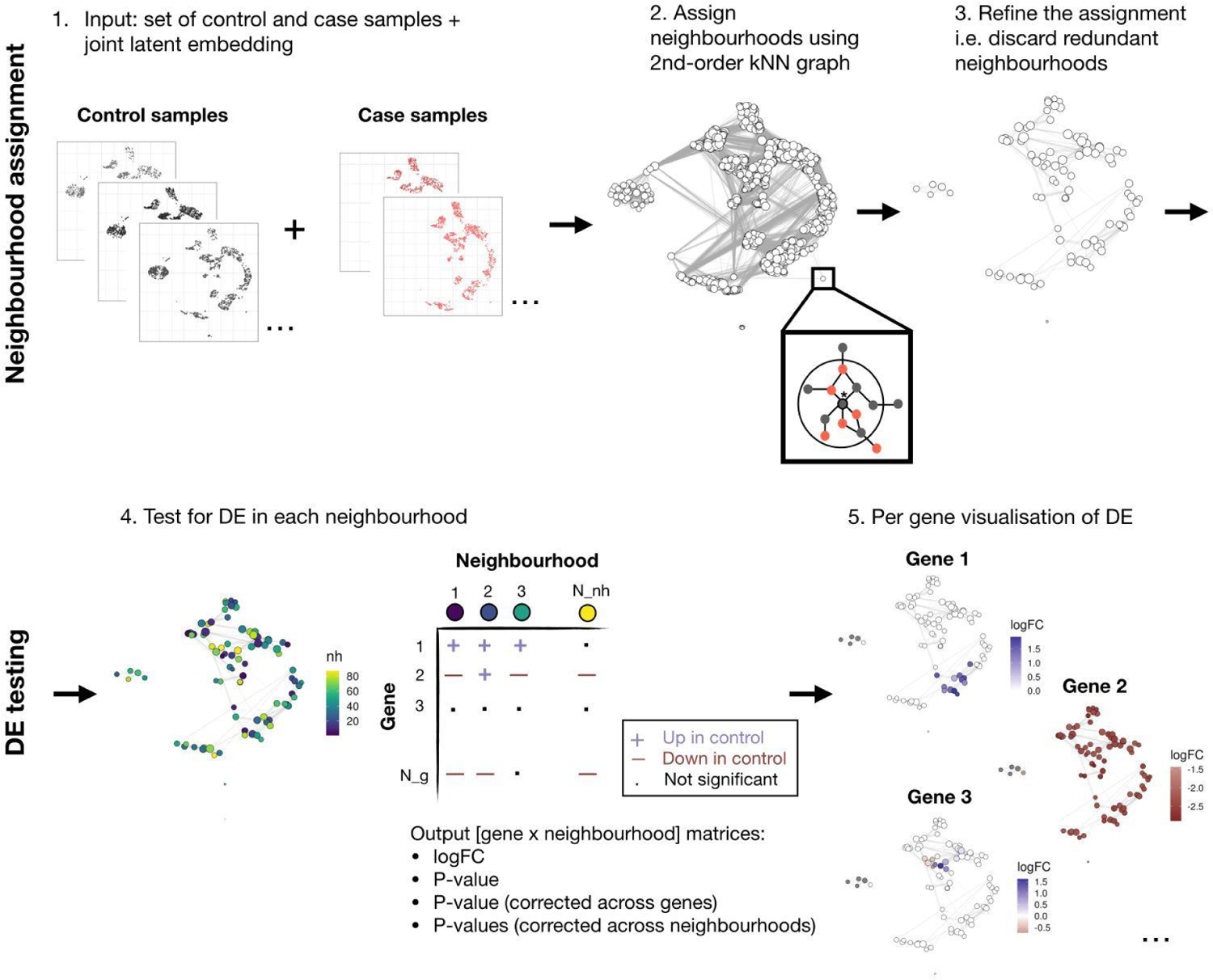
Schematic overview of the method. As input, the algorithm takes a set of samples with given labels (case or control) alongside a joint latent embedding (step 1). Next, we generate a graph recapitulating the distances between cells and define neighbourhoods (step 2) using the 2nd-order kNN graph (inset). We visualise the neighbourhood assignment with a ‘neighbourhood’ plot, in which each circle corresponds to the neighbourhood, and an edge between circles represents shared cells (the width of the edge corresponds to how many cells are shared between the neighbourhoods; for this cartoon, we discard all edges with weights less than 100). We then refine the neighbourhood assignment and discard redundant neighbourhoods (step 3). In step 4, we index our neighbourhoods, and then proceed with DE testing within each neighbourhood. As an output we return four [gene x neighbourhood] matrices, corresponding to logFC and statistics, raw and corrected either across genes or across neighbourhoods. Results for each gene can be visualised using neighbourhood plots (step 5), in which each neighbourhood (circle) is coloured with the estimated logFC if significant.

To construct a graph recapitulating distances between single cells we require count matrices and a pre-calculated latent embedding for all the samples from tested conditions (Fig. 1, step 1). Although the choice of the embedding/integration technique is left to the user, we note that supervised and unsupervised approaches result in neighbourhood assignments that differently impact DE detection. Specifically, we show that in supervised approaches, the input from case-specific variance in the embedding is smaller which is more suitable for DE detection within individual neighbourhoods (see in-depth discussion and supporting analysis in **Supplementary Note 1**).

A standard approach for constructing a graph representation is to use a kNN graph, in which each cell is connected with its *k* nearest neighbours in the latent space. One of the limitations of the kNN approach is its limited sensitivity to the local density of the data. In other words, if *k* is high enough, cells from rare cell types (i.e. less abundant than the chosen are prone to get connected (and therefore assigned to the same neighbourhoods) with transcriptionally similar but more abundant cell types (we discuss the importance of a sufficient number of cells for DE testing in **Supplementary Note 2**). To this end, several methods have been developed to improve on kNN graph representation to better recapitulate local density (Levine et al. 2015; Baran et al. 2019; Persad et al. 2022). These methods aim to identify phenotypically distinct, non-overlapping, fine-grained cell states. Motivated by this, to account for the local density we introduce a 2nd-order kNN graph representation in which a standard (hereafter referred to as a 1st-order) kNN graph is amended with additional edges between any 2 cells that share at least one neighbouring cell (Fig. 1, Step 2, inlet). As *k* increases, the average neighbourhood size for the 2nd-order kNN graph increases considerably faster than it does for the 1st-order, with a neighbourhood size in the low hundreds being achieved when *k* is between 20 and 30 (Supp. Fig. 1). Importantly, for cells from abundant cell types that contain many transcriptionally similar cells, the method will result in sufficiently large neighbourhood sizes. On the other hand, for rare cell types the corresponding neighbourhoods will be considerably more homogeneous, albeit smaller, compared to the 1st-order kNN graph, which is potentially more important for DE testing than neighbourhood size.

To quantitatively assess the detection differences between the assignments using either 1st- or 2nd-order kNN graphs, we used a chimeric mouse embryo dataset, in which tdTomato+ mouse embryonic stems cells containing a Tal1 knock out were injected into wild type blastocysts (Pijuan-Sala et al. 2019). We used an atlas of Wild Type (WT; i.e. non-chimeric) gastrulating mouse embryos as a “control” (henceforth referred to as WT), and the wild type cells from the chimeric embryos were used as the “case” (henceforth referred to as ChimeraWT). We assigned neighbourhoods using different *order* and *k* combinations while restricting *order-k* values such that the average neighbourhood size was within the low hundreds cell target (Supp. Fig. 2A). Since a certain degree of stochasticity, owing to the random selection of index cells, invariably impacts into any neighbourhood assignment, for each [*order-k*] combination (henceforth referred to as an assignment) we repeated the assignment 3 times.

**Figure 2.**
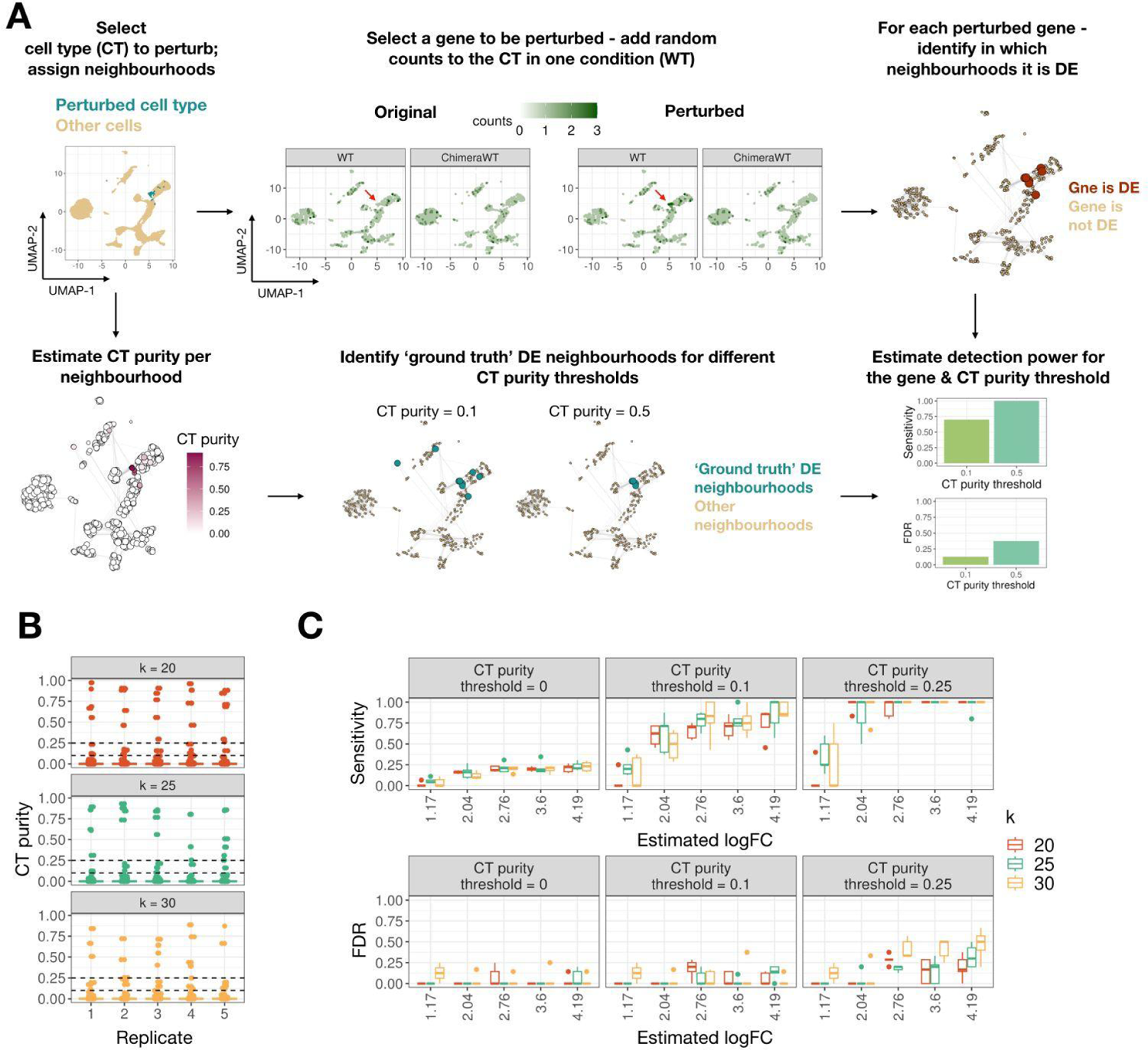
miloDE yields sensitive and precise detection in simulated data. A. Schematic representing how detection power is estimated for each gene and cell type purity threshold. Top left panel illustrates single-cell data, embedded in UMAP space, with highlighted cell type, in which we will alter the counts. Top middle panel represents the ‘in silico’ perturbation we introduce to the selected cells, and the top right panel represents neighbourhood assignment, followed by per neighbourhood quantification of whether the selected gene is identified as DE. Bottom left panel represents per neighbourhood quantification of the cell type ‘purity’. Bottom middle panels represent how ‘ground truth’ DE neighbourhoods are selected based on cell type purity threshold. Bottom right panel illustrates the final quantification of DE detection (i.e. sensitivity + FDR). B. Boxplots representing the distribution of cell type purity score across neighbourhoods for each neighbourhood assignment replicate and for *k* = 20, 25 and 30. Dashed lines correspond to selected cell type purity thresholds. C. Boxplots representing how sensitivity and FDR change with estimated logFC, *k* and cell type purity threshold. Each box represents data across 5 replicates for a single *k*.

Within each assignment, we calculated a per neighbourhood cell type purity score for three rare cell types (PGC, Definitive endoderm, and Floor plate; Supp. Fig. 2B). Cell type purity score for a cell type is defined as the percentage of cells from the neighbourhood that have the cell type label in question. To estimate whether assignments using the 2nd-order KNN graph contain neighbourhoods that are more ‘pure’ for rare cell types compared to the neighbourhoods from the 1st-order graph, for each assignment and each cell type we calculated the maximum cell type purity (across all neighbourhoods). We consistently observe higher cell type purity scores for the 2nd-order assignments across a wide range of average neighbourhood sizes (Supp. Fig. 2C).

Finally, we examined how cell type purity affects the sensitivity of DE detection. To do this, we synthetically introduced ‘ground truth DE’ by perturbing the expression of a small number of genes per cell type (see Methods). We then applied miloDE for each neighbourhood assignment and asked how the DE detection power (i.e. fraction of ‘perturbed’ genes that get detected as DE) depends on cell type purity as well as the absolute number of cells from the cell type. To do so, for each neighbourhood and each ‘perturbed’ cell type, we calculated the following measurements: cell type purity, the number of cells from the cell type, and DE detection power (Supp. Fig. 2D). As expected, DE detection power scales with both cell type purity and the number of cells from the cell type, but, more importantly, for the rarest cell type - PGCs - we achieve the highest detection power only for highly pure neighbourhoods, which are only present in the 2nd-order assignments. When we calculate the maximum DE detection power for each assignment, we observe a small but consistently higher detection power in PGC cells for the 2nd-order assignments, particularly for smaller neighbourhood sizes (Supp. Fig. 2E). Interestingly, the detection power for PGCs, on average, decreases with higher neighbourhood sizes, likely due to the fact that PGCs get ‘mixed’ with cells from other cell types. Overall, we conclude that 2nd-order assignments increase the power to detect DE in rare cell types.

In the original Milo approach, once the graph is constructed, the number of assigned neighbourhoods is controlled by the parameter *prop* - the initial proportion of cells selected as index cells (Methods). To decrease the computing time we want to minimise the number of tests (i.e. total number of neighbourhoods) while ensuring that all cells are assigned to at least one neighbourhood. Since it is unclear how to select the lowest *prop* while ensuring ‘complete coverage’, we introduce post hoc neighbourhood refinement (Fig. 1, step 3), in which we first assign a high enough number of neighbourhoods (we use *prop* = 0.2, but this can be increased by the user) to ensure complete coverage, followed by sorting the neighbourhoods in decreasing order of size and iteratively discarding neighbourhoods where all cells are included in previously assigned neighbourhoods (Methods). To test how post hoc filtering performs compared to selecting an optimal *prop*, we used the mouse gastrulation dataset introduced previously, constructed neighbourhoods with *prop* = 0.2 (i.e. use 20% of cells as index cells), and perform post hoc filtering. Based on the number of neighbourhoods after refinement, we then selected a considerably lower *prop* that resulted in a comparable number of neighbourhoods (without the refinement step). We observe that in the original, ‘unrefined’ neighbourhood assignments around 10% of cells are consistently unassigned to any neighbourhoods, whereas all cells are assigned in the refined assignment (Supp. Fig. 3).

**Figure 3.**
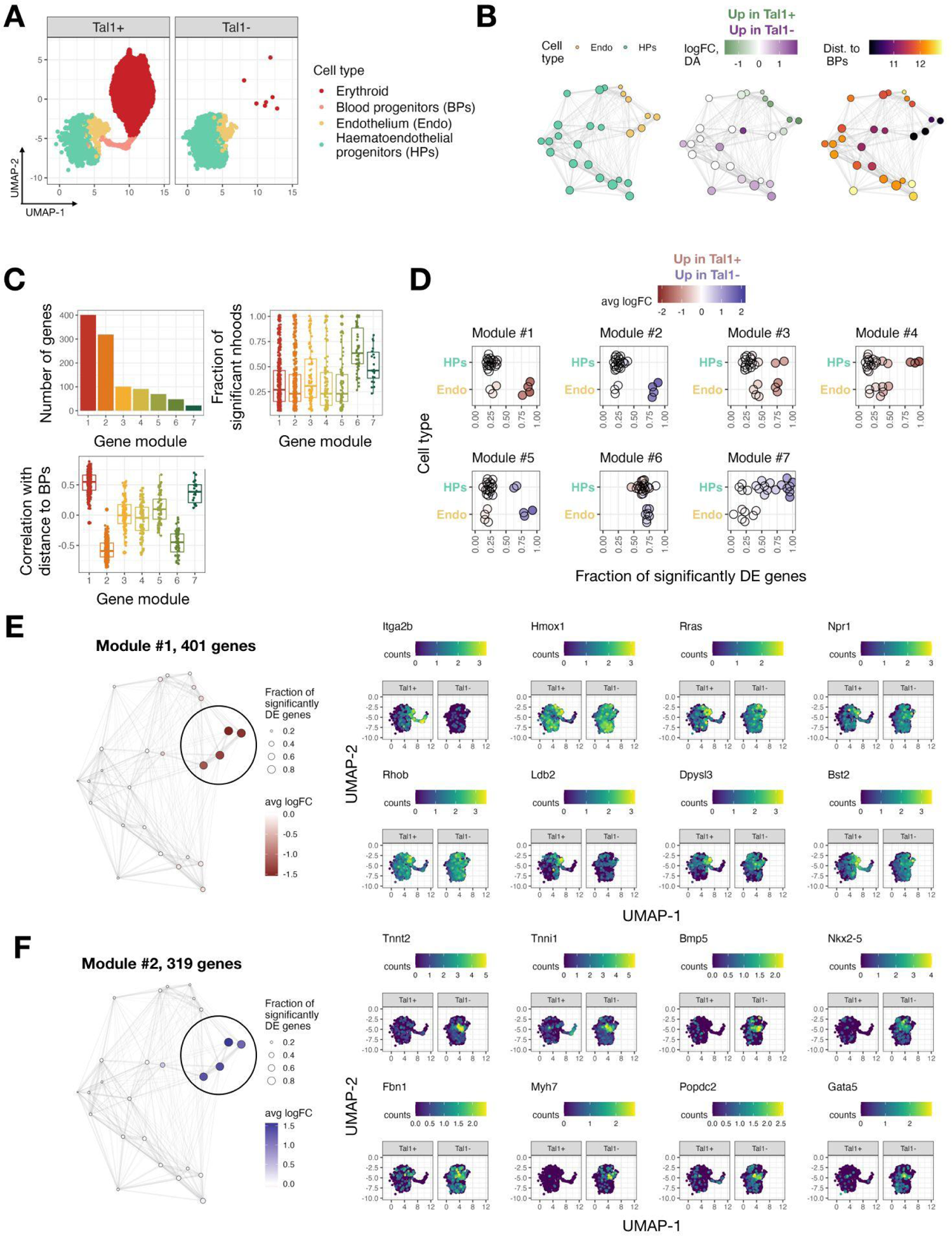
miloDE is suitable for continuous trajectories and recovers a transient transcriptional state of haemogenic endothelia. A. UMAP representing a manifold of chimera mouse embryos, colours correspond to the cell types contributing to blood lineage, and facets correspond to whether cells carry a knockout of Tal1. B. Neighbourhood plots covering haematoendothelial progenitors and endothelial cells. Each circle corresponds to the neighbourhood, and neighbourhoods are coloured by enriched cell type (left), differential abundance (middle) and distance to blood progenitor cells in PC space (right). C. Top left panel: Barplot representing how many genes are associated with each module. Top right panel: Distribution of fraction of significantly DE neighbourhoods for each module. Bottom panel: Distribution of correlation between logFC and distance to blood progenitors for each module. D. Jitter plot representing the relationship between gene modules and cell types. Each facet corresponds to one module, each point corresponds to one neighbourhood. X-axis corresponds to a fraction of genes from the gene module that are significantly DE in the corresponding neighbourhood, colour corresponds to average logFC. E. Left: Neighbourhood plot for the 1st gene module, size of the nodes correspond to fraction of genes from the gene module that are significantly DE in the corresponding neighbourhood, colour corresponds to average logFC. Right: UMAPs representing haematoendothelial progenitors and endothelial cells, coloured by representative genes from the 1st module that are associated with angiogenesis and lipoxygenase activity. F. Left: Neighbourhood plot for the 2nd gene module, size of the nodes correspond to fraction of genes from the gene module that are significantly DE in the corresponding neighbourhood, colour corresponds to average logFC. Right: UMAPs representing haematoendothelial progenitors and endothelial cells, coloured by representative genes from the 2nd module that are associated with cardiomyogenesis.

Following neighbourhood assignment, we test for differential expression within each neighbourhood (Fig. 1, steps 4-5). To decrease the burden of multiple testing correction, a standard practice in DE analysis is to discard genes that are lowly expressed in both case and control samples upstream of the testing (Bourgon, Gentleman, and Huber 2010). In miloDE, we provide a flexible framework to select genes that will be tested for DE, in which the user can regulate the minimum count value required for at least some samples (see Methods). In addition, akin to discarding ‘uninteresting’ genes, we also identify ‘uninteresting’ neighbourhoods where we anticipate no DE and discard these a priori (Methods, **Supplementary Note 3**). To test for DE within each neighbourhood, we use the edgeR framework - a scalable and highly performant DE detection method that utilizes a generalised linear model framework that allows the incorporation of covariates and complex experimental designs (Robinson, McCarthy, and Smyth 2010; Lun, Chen, and Smyth 2016). As an output, we return [gene x neighbourhood] matrices containing the estimated logFC and statistics indicating the significance of each comparison. We then perform multiple testing corrections in two directions: for each neighbourhood, we correct across all tested genes, and for each gene we correct across neighbourhoods, accounting for neighbourhood sizes (i.e. the number of cells per neighbourhood) and the overlap between neighbourhoods (see Methods). This dual correction scheme allows the identification of both DE-neighbourhoods for specific genes and DE-genes for specific neighbourhoods.

### miloDE enables sensitive and precise DE detection in simulated data

We next assessed the performance of miloDE in synthetically perturbed data. We used the same WT-ChimeraWT dataset introduced previously and perturbed the expression of 5 genes in one condition and in one selected cell type (henceforth ‘perturbed cell type’) (Methods, Fig. 2A). To ensure that the selected genes are not DE elsewhere on the manifold, we chose the genes from a pool of candidate genes that are not DE in any cell type. We assigned neighbourhoods and performed miloDE testing using *k* = 20, 25, 30 (2nd-order kNN), and for each *k* generated 5 replicates i.e. independent neighbourhood assignments (Fig. 2B). Next, for each neighbourhood assignment, we calculated the per neighbourhood cell type purity i.e. how many cells from a neighbourhood have the perturbed cell type label. We define neighbourhoods as ‘ground truth’ DE if cell type purity for these neighbourhoods exceeds a certain cell type purity threshold for the perturbed cell type. Finally, for any ‘perturbed’ gene and any cell type purity threshold, we can calculate the detection power for this gene (i.e. sensitivity and FDR) based on the overlap between ‘ground truth’ DE neighbourhoods and neighbourhoods in which the gene is identified as DE. We then assessed how sensitivity and FDR change as a function of the cell type purity threshold (Fig. 2C). Consistent with the expectation that the probability of detecting DE in a neighbourhood scales with both the fraction of ‘altered’ cells as well as the effect size, we observe that sensitivity and FDR increase as a function of both cell type purity threshold and estimated logFC. We observe reasonably high sensitivity when using a cell type purity threshold of 0.1 (i.e. all neighbourhoods in which at least 10% of cells belong to the perturbed cell type are flagged as ‘ground truth’ DE) which suggests that even a fraction of perturbed cells in one condition is sufficient for a test result to be flagged as significant. Importantly, we report reasonably high sensitivity with well-controlled FDR, across a wide range of cell type purity thresholds and estimated effect sizes. Moreover, these trends are robust across different levels of *k*. Overall, we conclude that miloDE provides robust, sensitive, and precise DE detection across a wide range of effect sizes.

### Identification of haemogenic endothelial-like cells undergoing ectopic cardiomyogenesis in the absence of TAL1

Having established the good performance of miloDE, we applied it in the context of continuous developmental trajectories, where discrete clustering and cell type annotation is suboptimal. Tal1 (SCL) is a DNA-binding transcription factor that plays a key role in haematopoiesis, with mouse embryos that carry a double knockout of *Tal1* dying around embryonic day E9.5 from severe anaemia (Shivdasani, Mayer, and Orkin 1995). Previously, the molecular function of Tal1 has been investigated using chimeric mouse embryos where *Tal1*-/- mouse embryonic stem cells are injected into wild-type blastocysts; the wild type blastomeres are able to generate blood cells, meaning that the cell-intrinsic impact of Tal1 can be effectively studied (Robb et al. 1996; Pijuan-Sala et al. 2019). As shown previously, in *Tal1*-chimeras the *Tal1* mutant cells are depleted of erythroid cells (Fig. 3A, Supp. Fig. 4A). Additionally, a laborious annotation of endothelial cells revealed that mutant cells that transcriptionally resemble haemogenic endothelia (which in the wild-type environment would have contributed to the second wave of hematopoiesis) do not express haemogenic markers and instead show signs of cardiomyogenesis.

**Figure 4.**
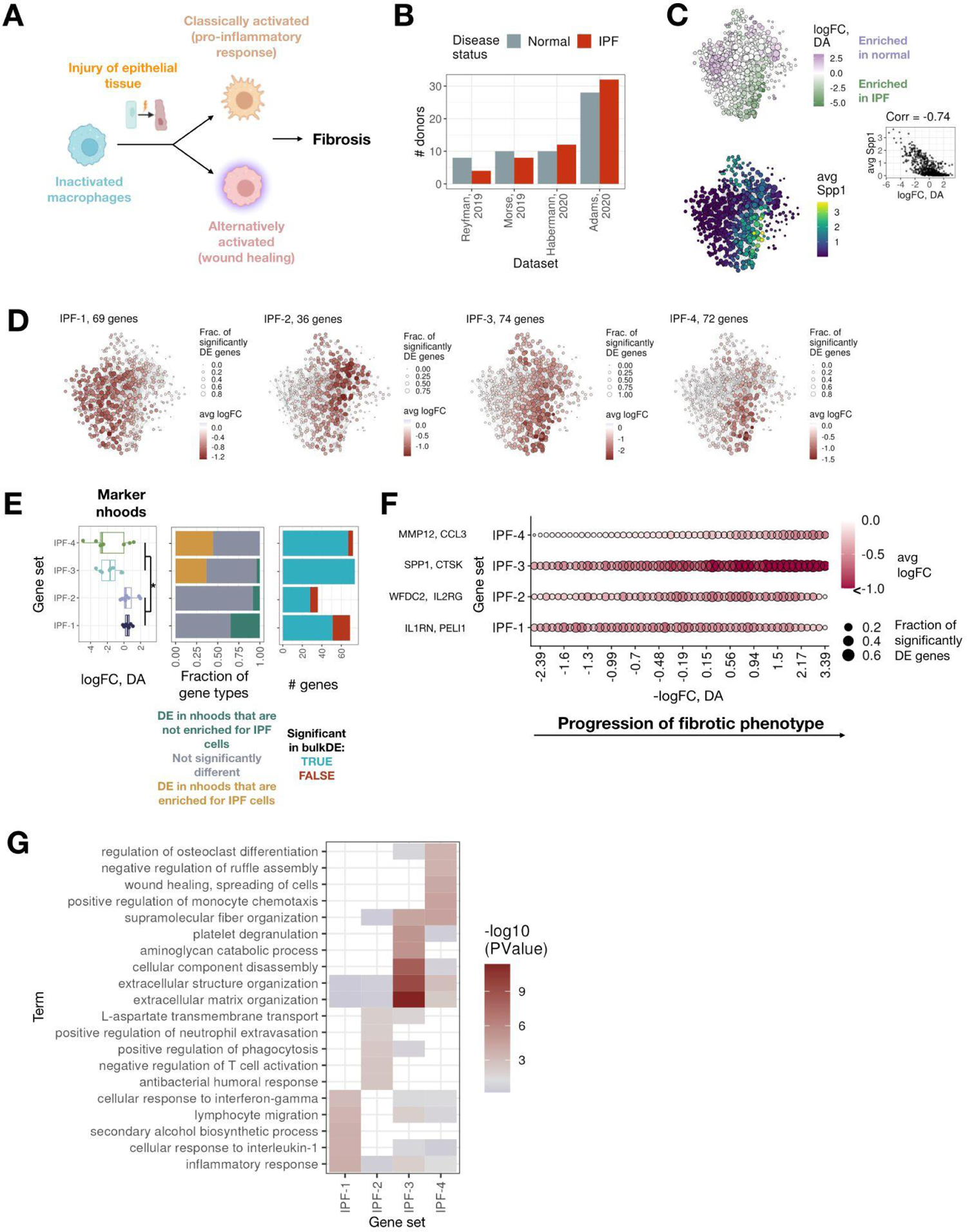
miloDE enables sufficient resolution to discover macrophage-specific gene sets, specific for different points of the IPF progression. A. Schematic cartoon (based on a figure from Zhang et. al. 2018) representing the phenotypical heterogeneity of activated macrophages upon the response to injuries in epithelial tissues. B. Barplot representing donor composition across different datasets and conditions. C. Top left panel: The neighbourhood plot covering macrophages in both healthy and IPF patients, each node is coloured by estimated differential abundance. Bottom left panel: The neighbourhood plot covering macrophages in both healthy and IPF patients, each node is coloured by average Spp1 expression. Middle right panel: High negative correlation (across neighbourhoods) between differential abundance and average Spp1 expression reflects enrichment of Spp1 fibrotic macrophages in IPF patients. For the neighbourhood plots, we discard all edges with weights less than 100. D. Neighbourhood plots representing transcriptional profiles for IPF-upregulated gene sets. For each gene set, size of the nodes correspond to the fraction of genes from the gene set that are significantly DE in the corresponding neighbourhood, and colour represents average logFC. E. Left panel: Boxplots representing the distribution across ‘marker’ neighbourhoods for each cluster (y-axis) with the respect to logFC of the differential abundance (x-axis). Asterisk signifies a significant difference between first 2 and last 2 gene sets (p-value < 0.05, Wilcoxon rank test). Middle panel: Barplot representing a composition of different gene categories across gene sets. Right panel: Barplots representing whether genes are detected as DE in the bulk (across whole cell type) testing. F. Dotplot representing, for each gene set, its aggregated DE pattern along the fibrotic phenotypical progression (fibrotic progression is estimated from logFC of differential abundance testing). Colour of the dots corresponds to aggregated average logFC and the size of the dots corresponds to the average fraction of genes from the gene set that are significantly DE in the corresponding neighbourhood. G. A heatmap representing functional annotation across gene sets. Y-axis corresponds to enriched gene ontologies, x-axis corresponds to the gene sets, and colour corresponds to the significance of the GO enrichment for the corresponding cluster.

To investigate if miloDE could provide further insight into this biological process, we first applied it to the whole manifold (i.e. all cell types) to identify which cell types show signs of extensive transcriptional changes upon the perturbation. To rank cell types by ‘degree of perturbation’, for each neighbourhood, we calculated the number of DE genes as well as the number of DE genes that are highly specific to this neighbourhood (see Methods), followed by grouping neighbourhoods by the cell type they primarily contained (Supp. Fig. 4B, C). Reassuringly, neighbourhoods that are enriched for cells directly contributing to the blood lineage (i.e. endothelial and haematoendothelial progenitors) rank highest for both metrics (with the exception of one neighbourhood, containing a mixture of rare cell types that are only present in one condition, Supp. Fig. 4D), with all other cell types showing a considerably lower degree of perturbation. This observation supports the expectation that while endothelia and haematoendothelial progenitors are present in both conditions in similar quantities, the absence of Tal1 still results in transcriptional changes that we can identify and characterise using miloDE.

To systematically assess how hematopoiesis is disrupted in cells contributing to the blood lineage, we next applied miloDE only to cells annotated as haemoendothelial progenitors and endothelia (Fig. 3A). To characterise DE-patterns, we calculated various statistics on the assigned neighbourhoods, including cell type enrichment, differential abundance and average distance to blood progenitors (Methods, Fig. 3B). To further explore DE patterns within the selected cells, we applied the WGCNA framework on logFC vectors and retrieved transcriptional modules of co-perturbation - sets of genes that show similar magnitude of DE across the neighbourhoods, and are thus likely to be co-regulated (Langfelder and Horvath 2008; Langfelder et al. 2011; Feregrino and Tschopp 2022, Methods). We identified 6 co-regulated modules, containing different numbers of genes (Fig. 3C). Most modules contain genes that are differentially expressed in coherent sub-regions of the manifold (Fig. 3C, Supp. Fig.5). Additionally, for several modules, the associated regions of DE (i.e. neighbourhoods in which the majority of the genes from the module are DE) contain neighbourhoods associated with both endothelia and haematoendothelial progenitors (Fig. 3D). This likely reflects the limitation of using discrete clustering approaches to summarise the continuous trajectory of endothelial maturation.

Next, we focused on the first two modules, which are ‘complementary’ to one another. Specifically, for each gene we calculated the correlation between its logFC and distance to blood progenitors (across neighbourhoods), and based on it, we suggest that module one contains genes that are down-regulated (in *Tal1-* cells) and module two contains genes that are up-regulated in the neighbourhoods that are most proximal to blood progenitors. Importantly, their ‘location’ in the manifold suggests that they may be enriched for cells that give rise to blood lineage (Methods, Fig.3B-D). Consistent with this, we find that the first module contains the hematopoietic marker *Itga2b* (Dumon et al. 2012; McGrath et al. 2015) as well as genes associated with angiogenesis and vasculature development (*Rras*, *Hmox1*, *Npr1*, and others; Fig. 3E, Supp. Table 1) and the regulation of cell migration (*Rhob*, *Dpysl3*, *Ldb2*, *Bst2*, and others; Fig. 3E, Supp. Table 1), which is characteristic of the endothelial to hematopoietic transition (Ottersbach 2019). The second module, containing genes that are upregulated in *Tal1*-cells, is heavily enriched for genes associated with cardiomyogenesis (*Tnnt2*, *Tnni1*, *Bmp5*, *Nkx2-5*, *Gata5,* and others; Fig. 3F, Supp. Table 1). This is consistent with the observations from the original study as well as previous reports suggesting an alternative, cardiac, cell fate specification in mesodermal cells that lack *Tal1* (Org et al. 2015; Scialdone et al. 2016; Chagraoui et al. 2018; Pijuan-Sala et al. 2019). The functional annotation of the first two modules, together with the fact that associated neighbourhoods are transcriptionally ‘close’ to blood progenitors, strongly suggests that these neighbourhoods consist of haemogenic endothelia that in normal circumstances give rise to blood cells during the second wave of hematopoiesis. Intriguingly, within this region, we also observe sign difference in differential abundance, with the neighbourhoods most proximal to blood progenitors being enriched for Tal1+ cells. This pattern likely reflects the inability of Tal1-endothelial cells to commit to a haemogenic identity, followed by the block in their development and acquisition of cardiac fate. Combined, our findings confirm the sensitivity of miloDE to identify transient cell states in continuous manifolds, within and across individual cell types.

### miloDE reconstructs a functionally meaningful timeline of macrophage activation during Idiopathic Pulmonary Fibrosis (IPF)

To showcase the power of miloDE in the context of complex diseases and datasets, we next applied it to study transcriptional changes that arise in Idiopathic Pulmonary Fibrosis (IPF). IPF is a severe, irreversible lung condition that initiates with persistent inflammation of epithelial cells, which in turn triggers an inflammatory response resulting in constant wound healing and scarring of the epithelial tissues of the lungs (fibrosis) (Lederer and Martinez 2018).

At the cell type level, macrophages are one of the key players in IPF. Macrophages are phenotypically plastic cells, with their transcriptional make-up being highly dependent on their environment and the external stimulant. Consistent with this, their function and role in fibrosis changes with disease progression (Alber et al. 2012; Zhang et al. 2018; Ogawa et al. 2021). At the early stages of the disease, characterized by the inflammation of epithelial cells lining the airways, ‘classically activated’ macrophages are observed (Fig. 4A). As a direct response to the inflammation, these macrophages are activated by IFN-gamma or lipopolysaccharides and produce pro-inflammatory cytokines. In turn, a persistent inflammatory response triggers aberrant wound healing and fibrotic remodelling, associated with ‘alternatively activated’ macrophages that can be activated by IL-4/13.

By definition, DE approaches applied to a specific cell type will return, for each gene, a single logFC, rendering it impossible to characterise different continuous patterns of DE changes within the cell types. Importantly, at cluster resolution, it is also impossible to distinguish DE caused by sub-cell type differential abundance, from disease-associated DE. On a molecular level, one axis of macrophage variability in IPF is associated with the expression of the fibrotic marker Spp1 (Morse et al. 2019; Reyfman et al. 2019; Adams et al. 2020). The abundance of an Spp1-high subpopulation has been repeatedly observed in IPF patients, however, Spp1-high macrophages also exist in healthy patients (albeit at a lower abundance and absolute Spp1 expression). This expected sub-cell type compositional biases between normal and IPF patients might lead to false positive detection of DE genes ‘marking’ transcriptionally different subpopulations in macrophages. Consequently, a cluster-free approach is necessary to study subtle transcriptional changes that arise in this population of cells during disease progression.

To this end, we analysed an existing lung atlas available within Azimuth, considering four datasets that contain cells from healthy and IPF donors (Fig. 4B). In each of these, we only considered cells annotated as macrophages. First, we assessed DA across neighbourhoods within the macrophage manifold, observing (as expected) differences in abundance that are highly correlated with average Spp1 expression (Fig. 4C), suggesting that the logFC of DA can be used as a proxy for disease progression.

We then applied miloDE and characterized different DE patterns by applying Louvain clustering to group genes upregulated in at least 10% of neighbourhoods into gene sets, using their logFC values across neighbourhoods as a feature vector ((Blondel et al. 2008), Methods). This analysis identified 4 gene sets (Fig. 4D); interestingly, neighbourhoods strongly associated with each set were enriched for specific stages of disease progression (Fig. 4E, left panel). Specifically, ‘neighbourhood markers’ for the first two gene sets have logFC-DA around 0 (i.e. neighbourhoods that contain comparable numbers of IPF and healthy cells), whereas the last two gene sets are marked by neighbourhoods that mostly show negative logFC-DA (i.e. enriched for IPF cells). Moreover, even though we focused our analysis on genes that are upregulated in at least 10% of the neighbourhoods, ∼12% of the genes that we used for Louvain clustering were identified as not significant when DE was performed on the pseudo-bulk (p-value > 0.05, DE across all macrophages) level and this under-calling is particularly prominent in the ‘earlier’ gene sets (Fig. 4E, right panel). Among the genes that were not detected at the bulk level are members of the immunoglobulin family (*Iglc2*, *Ighg1*, *Ighg4*), genes associated with the defence response against bacteria (*Plac8*, *Defb1*), and regulation of B cell activation (*Peli1*).

To further characterise gene sets, we next assigned each gene into one of three groups: DE in neighbourhoods that are enriched for IPF cells, DE in neighbourhoods that have both IPF and healthy cells in comparable amounts, and DE in both groups of neighbourhoods (Fig. 4E, middle panel; Methods). We observe that sets that are associated with neighbourhoods that are enriched for IPF cells (i.e. IPF-3 and IPF-4) contain genes that are identified as DE predominantly in IPF-enriched regions of the manifold, whereas sets IPF-1 and IPF-2 contain genes that are DE in neighbourhoods containing similar number of normal and IPF cells (i.e. not DA neighbourhoods).

To systematically assess the relevance of each gene set along the phenotypic progression of fibrotic macrophages, we used the logFC-DA to combine neighbourhoods into 50 bins, and calculated the average fraction of significant genes and average logFC (Fig. 4F). The first 2 sets contain genes that are mainly DE (i.e. with a high fraction of genes being DE) at the early to mid stages of the disease (Fig. 4F; Supp. Fig. 6). On the other hand, the last 2 gene sets are mainly DE at the late stages of the disease. Specifically, IPF-3 contains genes that are systematically DE across all macrophages, with the effect size increasing with disease progression. We note that most of the known macrophage markers such as *Spp1*, *Csf1*, *Sparc*, *Ctsk*, *Gpc4,* and *Mmp9* (Adams et al. 2020) fall into this gene set. Finally, IPF-4 contains genes that are DE specifically in the later stages of the disease.

When we perform functional annotation of the gene sets (Fig. 4G), we observe strong agreement with the previously suggested functional ‘timeline’ of macrophage activation. Specifically, the IPF-1 gene set is significantly enriched for inflammatory response genes and, in particular with genes associated with cellular response to interleukin-1 (*Ccl18, 23, 24; Peli1*, *Il1rn*), coherent with the observation that classically activated macrophages release IL-1beta chemokines (Murray and Wynn 2011; Zhang et al. 2018). Intriguingly, *Il1rn* is a potential therapeutic target of IPF (Borthwick 2016), and in IPF-1 we also see the enrichment of genes associated with the ERK pathway (*Ccl18, 23, 24; Pdgfd*), inhibition of which prevents the progression of established fibrosis (Madala et al. 2012). We suggest that miloDE provides a powerful exploratory framework that can aid the identification of subtle changes in gene expression that arise early in disease progression. Genes from IPF-2 are associated with an ‘initial’ defense response against pathogens (*Defb1, Wfdc2, Il2Rg, Fcgr2b*), consitent with the observation that pathogenic bacteria is one of the drivers of IPF progression (Yang et al. 2015; Fastrès et al. 2017). In contrast to the first 2 gene sets, IPF-3 is characterized by the hallmarks of a more advanced state of fibrosis such as extracellular matrix deposition (*Spp1*, *Sparc*, *Fn1*, *Ctsk*, *Mmp7,9,10,* etc), cellular disassembly, and platelet degranulation (*Timp3*, *Cd9*, *Ppbp*, *A2M*). Finally, IPF-4 contains genes marking pathogenic, late-stage fibrosis such as wound healing (*Mmp12*, *Rhoc*, *Arhgap24*) and collagen fibril organisation (*Col4a2*, *Col6a2, Col8a2*). Combined, we illustrate that miloDE enables the discovery of subtle differences in gene regulation, with potentially important implications for how scRNA-seq data can be used to study the early stages of disease.

## Discussion

The amount and diversity of scRNA-seq data is revealing ever more complex and subtle patterns that characterise processes ranging from normal development through to the onset and progression of disease. Whilst broadly applied clustering strategies can provide high-level annotation of the heterogeneity within individual organs and tissues, their discrete nature is suboptimal for analysing more granular changes in expression, including in the context of continuous trajectories. miloDE addresses this challenge in the context of testing for differential expression. Leveraging overlapping cell neighbourhoods overcomes the tedious, frequently inaccurate, and time-consuming step of clustering and cell type annotation. Consequently, miloDE allows an in-depth characterisation of various DE patterns, both across the whole dataset and within individual cell types. Given the rapid advances of extensive cohort atlas studies and large-scale CRISPR screens, miloDE is a timely computational tool, which successfully leverages a large number of replicates in order to enable sensitive and specific DE detection, while accounting for complex experimental designs.

While miloDE approximates gene regulation as a more continuous function of the manifold, it is important to acknowledge that it leverages the same principle of cell grouping as standard per cell type approaches, and therefore it is prone to similar pitfalls. To enable sensitive DE detection, we require that neighbourhoods exceed a certain size, which in turn can compromise the homogeneity of the cells in the neighbourhoods. Within miloDE we address this by introducing a 2nd-order kNN graph representation that provides an improved estimation for the local graph density. However, given the target number of cells per neighbourhood, it is impossible to guarantee that transcriptionally distinct cells will not get assigned to the same neighbourhood. This challenge can be partially mitigated by focusing on genes with strong effect size and by decreasing *k*.

Another important limitation that stems from using overlapping neighbourhoods is the potential propagation of false positives. Specifically, while this approach overcomes limitations associated with discrete clustering, single cell(s) with an altered transcriptional makeup can dominate DE results in multiple tested groups. In other words, if a single cell has extremely high (potentially sporadic) expression of a certain gene, some of the tested groups including this cell might be indicated as DE for this sporadic gene. Importantly, the issue scales with the increase in *k* (which results in a higher degree of overlap between the neighbourhoods), but it is also partially relieved by the neighbourhood refinement step and can further be controlled by using a larger number of biological replicates. Another inevitable source of false positives is the extensive number of tests we perform - while the relative number of false positives (after multiple testing correction) is likely to be low (Fig. 2), the absolute number can still be substantial. Importantly, while individual gene-neighbourhood combinations might result in false positive identification of DE and will require a manual examination and orthogonal verification, we suggest that the true exploratory power of miloDE lies in its ability to identify subtle and local patterns of co-regulated genes (thus minimising the input from random FPs).

In this work, we adapted two readily available approaches for identifying DE-patterns - WGCNA and Louvain clustering. While they both have proved useful and have aided in biological discoveries, it is important to highlight their limitations. The WGCNA-based approach is an exclusive process meaning that only a fraction of genes will be assigned to any module, and therefore it is possible that interesting DE patterns will be excluded. In addition, the WGCNA-adapted approach to detect co-regulated gene modules is highly sensitive to input data, thus impeding the interpretation of the modules. On the other hand, the Louvain-adapted approach is inclusive of all input genes. However, clustering can be driven by a handful of neighbourhoods, with the rest of the manifold being incoherently DE within individual gene clusters, thus rendering the characterisation of the clusters challenging. Nonetheless, the combination of the suite of analysis methods allows for a flexible and tailored interpretation of the miloDE output.

Finally, moving forward, we suggest that in parallel with the emergence of novel cluster-free computational approaches for comparative analysis, it is crucial to contemplate how we will handle and interpret such complex outputs. One, nearly philosophical, conundrum is the decoupling between differential abundance and expression. While a mild change in expression for a certain gene in a certain transcriptional region can be detected as DE, it is possible that a stronger change in the expression will lead to the transition between transcriptional changes (i.e. DA). However, this depends on whether the gene in question contributed to the embedding in the first place. miloDE is uniquely able to analyse DE and DA in parallel by combining the original Milo approach with miloDE. In sum, we believe that miloDE is a good stepping stone toward continuous comparative analysis, which is where the field of single-cell analysis is heading.

## Methods

### Code availability

Scripts to generate data and to perform the above analysis are available at https://github.com/MarioniLab/miloDE_analysis.

### miloDE pipeline

miloDE is a cluster-free Differential Expression (DE) framework that leverages a graph representation of single-cell data. In the first step of the pipeline, miloDE performs the assignment of cells to (overlapping) neighbourhoods; each neighbourhood contains neighbouring cells, estimated from the graph representation. Once neighbourhoods are assigned, DE testing is carried out for each neighbourhood individually, and for each neighbourhood and gene that was tested in this neighbourhood, we return a logFC and p-value. Once the testing is concluded, we perform multiple testing correction in two directions: for each gene, across tested neighbourhoods, and for each neighbourhood, across tested genes. The final output consists of 4 [gene x neighbourhood] matrices, containing logFC estimated, uncorrected p-value, corrected p-value across neighbourhoods, and corrected p-value across genes.

Below we provide more details for each step:

1. *Neighbourhood assignment*.
  1.1. Input. As input, the algorithm takes a SingleCellExperiment object (Amezquita et al. 2020) containing a count matrix for all the samples combined. We require that the `colData` slot contains entries that identify sample IDs (used as replicates in DE testing) and conditions to be tested. We also require that a pre-calculated joint latent embedding is provided in the `reducedDim` slot.
  1.2. Graph construction and neighbourhood assignment. We use 1st- or 2nd-order kNN graphs to represent transcriptional relationships between cells and assign neighbourhoods (the order of the graph is specified by the user, default is 2). We define a 1st-order kNN graph as a standard kNN-graph, and the 2nd-order kNN graph as a 1st-order kNN graph augmented with edges between cells if they share at least one cell within their neighbours. Neighbourhoods with the centre cell *c* are defined as all cells (including *c*) that are connected with it by the edge. In order to define neighbourhoods on a graph representation, we first select index cells (or cells that we will use as a neighbourhood centre in order to assign neighbourhoods). Since the computational complexity of identifying index cells scales with the average density of the graph, and the 2nd-order graphs on average are anticipated to be dense, we perform graph assignment and search for the index cells in parallel. Specifically:

1. We construct a 1st-order kNN graph, with *k* = min(50, *k*), where *k* is an input of the algorithm and has to be provided by the user.
2. We then use this graph to select index cells using the waypoint sampling algorithm used in Milo (Csardi, Nepusz, and Others 2006; Gut et al. 2015; Dann et al. 2021).
3. Once index cells are selected, we recalculate the 1st-order kNN graph if *k* > 50.
4. If *order* = 2, we construct the 2nd-order kNN graph by adding edges between vertices if they share at least one vertex as their neighbour.
5. Once the graph is constructed and index cells are selected, for each index cell we assign all its neighbours into a single neighbourhood. As an output of this step, we return the [cell x neighbourhood] matrix with boolean identifiers of cell inclusion in the neighbourhood.
  1.3. Neighbourhood refinement. Graph refinement is an optional, but recommended step, during which we identify neighbourhoods that can be discarded without any cells being unassigned to at least one remaining neighbourhood. The graph refinement process represents a case of the NP-hard ‘set cover problem’. Accordingly, we use a heuristic greedy implementation to solve the cover problem provided by the `Rcpp::greedySetCover` function (Eddelbuettel and Francois 2011; Eddelbuettel n.d.). Within this implementation, sets are first sorted in decreasing order of their power (i.e. how many elements they contain), and then iteratively a set is included if it includes at least one element that does not belong to any previously included sets. In the context of miloDE neighbourhood refinement, we first assign an excessive number of neighbourhoods to ensure that all cells are ‘covered’ at least once; we use neighbourhoods as sets and cells as elements.
2. DE testing.
  2.1. Selection of neighbourhoods to be tested. As an optional step of the algorithm, we perform a selection of neighbourhoods that show signs of ‘expression shifts’ and therefore are recommended to be tested for DE. Discarding ‘unperturbed’ (i.e. with no signs of ‘expression shifts’) neighbourhoods is computationally beneficial for downstream steps of the pipeline as well as facilitating the burden of multiple testing correction. To identify potentially ‘perturbed’ neighbourhoods, we adapt Augur, an approach that was originally designed to rank cell types by the degree of their perturbation in a specified condition (Skinnider et al. 2021). Specifically, for each cell type, Augur builds a Random Forest classifier that predicts the condition to which a cell belongs, and for each cell type returns AUCs. To adapt Augur in the context of miloDE, for each neighbourhood we return the AUC from the classifiers implemented by Augur. The user can then select their own AUC threshold to decide which neighbourhoods should be supplied further for DE testing. Note that our simulations show that in the presence of unbalanced batch effects nearly all neighbourhoods, regardless of whether simulations contained DE genes or not, had AUC > 0.5 (which is a recommended default value for the AUC threshold). Since this step can be time-consuming, we recommend including it in cases where the batch effect is anticipated to be minimal.
  2.2. Selection of genes to be tested. We provide a framework to select genes to be tested. To do so, we employ `edgeR::filterByExpr` to determine which genes have sufficient counts to be considered for DE testing. The user can tune the minimum count required for at least some samples by changing the `min_count` parameter (default is 3). This procedure is performed for each neighbourhood separately, which might result in different sets of genes being tested within each neighbourhood. Note that `min_count` can be set to 0, and in this case no gene selection will performed.
  2.3. DE testing within neighbourhood. To carry out DE testing, we use quasi-likelihood testing from `edgeR,` which was originally designed to perform DE testing on bulk RNA-seq. Specifically, to perform DE within each neighbourhood, the algorithm proceeds as following:

- We filter out all samples that contain less than `min_n_cells_per_sample` parameter, specified by the user (default is 3).
- For each sample present in the neighbourhood (i.e. biological replicate), we aggregate counts to create pseudo-bulks that mimic bulk RNA-seq data.
- We select genes as described above, and using total sum counts across selected genes as a proxy for library size, calculate offset factors (`edgeR::calcNormFactors`) in order to correct for compositional biases.
- We use the experimental design provided by the user. We allow the incorporation of covariates that are stored in the metadata of the counts matrix.
- We estimate the Negative Binomial dispersions as a function of gene abundance (`edgeR::estimateDisp`).
- We estimate quasi-likelihood dispersions (`edgeR::glmQLFit`). The quasi-likelihood dispersion models variability of the per gene estimated variances.
- We perform DE testing using a generalised linear model (`edgeR::glmQLFTest)`. As an output, we get a table, where rows correspond to the tested genes, and columns contain estimated logFC, p-value, and FDR (i.e. p-values corrected across tested genes).
  2.4. Multiple testing correction across neighbourhoods. Once tests for each neighbourhood are performed, for each gene we correct calculated p-values across the tested neighbourhoods. To do so, we adapt the spatial correction approach originally introduced in CYDAR (Lun, Richard, and Marioni 2017) and further developed in Milo (Dann et al. 2021). Specifically, we leverage the graph representation used to assign cells to neighbourhoods, and apply the weighted Benjamini-Hochberg correction approach, where p-values for each neighbourhood are weighted by the reciprocal of their local density. As a proxy for the local density for each neighbourhood, we use a weighted sum across all cells in the neighbourhood, where weights are calculated as the number of neighbourhoods to which a cell belongs. Note, that graph-based estimation for neighbourhood density was developed after the original Milo publication, and available in the Bioconductor version 3.16 (`BiocManager::install(version = “3.16”)`). Finally, we only perform the correction across the neighbourhoods for which testing for the gene was carried out. For neighbourhoods for which testing was not performed, we return NaN.

### Analysis of the effect of embedding approaches on miloDE performance

1. *Dataset.* We use mouse gastrulation scRNA-seq data from (Pijuan-Sala et al. 2019); data downloaded using `MouseGastrulationData()` package (Griffiths and Lun, 2022). As WT cells, we use all samples from the E8.5 developmental stage. As ChimeraWT cells, we use tD-tomato negative cells from Tal1-chimera mouse embryos (which are also from developmental stage E8.5). Next, we concatenated counts from both experiments, and embeddings were calculated on log-normalised counts (in batch aware manner, using batchelor::multiBatchNorm (Haghverdi et al. 2018)).
2. *Embeddings.* We analyse several embedding approaches, including both unsupervised (MNN (Haghverdi et al. 2018)) and supervised (Azimuth (Hao et al. 2021), reference-projected MNN, and scArches (Lotfollahi et al. 2022)). All approaches perform batch-aware embedding, and sample ID was used as the batch. For all embeddings, except scArches, we calculate a 30-dimensional latent space. For scArches we use the provided default (10). Below we introduce more detailed descriptions for each approach.

- MNN is an unsupervised approach that performs batch-aware PCA correction followed by MNN correction. The MNN-based latent space is sensitive to the selected genes for which we perform PCA. Accordingly, we select 3000 highly variable genes using `scran::getTopHVGs` (Lun, McCarthy, and Marioni 2016), and the selection is performed using either only WT data; only ChimeraWT data; WT + ChimeraWT data.
- Reference-projected MNN is a supervised variation of the original MNN. Specifically, we first perform batch-aware PCA on control data using `batchelor::multiBatchPCA`. As an output, we get batch-corrected PCs for the control data as well as the matrix of rotation vectors that we can apply to calculate PCA on scaled case data. Finally, we concatenate the calculated PCs and perform MNN-correction using `batchelor::reducedMNN`.
- Azimuth integration follows the pipeline for the reference-based mapping introduced by (Hao et al. 2021) and employed by the Azimuth web application for reference-based single-cell analysis (https://azimuth.hubmapconsortium.org/). To perform integration, we follow the provided by the Azimuth mapping pipeline.
- scArches integration follows the pipeline from (Lotfollahi et al. 2022).
3. *Chimera-specific DE genes.* To identify chimera-specific genes that are systematically DE across all cell types, we performed DE testing for each cell type for which we have at least 100 cells and selected genes that are DE (FDR < 0.05) in all tested cell types. We then ordered all genes according to the average logFC, and selected the top 3 genes with the highest absolute average logFC (*Rpgrip1*, *Slc24a5*, *Cbx3*).
4. *Neighbourhood assignment and estimation of neighbourhood homogeneity.* For each embedding, we generated a kNN graph (*k* = 100, *order* = 1) and assigned cells to neighbourhoods. We then discarded redundant neighbourhoods as described above. To assess the homogeneity of the neighbourhoods, for each embedding and each chimera-specific DE gene, we calculated the standard deviation of log-normalised counts within each neighbourhood, separately across WT and ChimeraWT cells.
5. *Estimation of logFC for chimera-specific DE genes*. To estimate logFC distribution for each embedding and each chimera-specific DE gene, we performed miloDE for each embedding (neighbourhood assignment based on kNN graph, *k* = 100) and aggregated logFC estimates across all neighbourhoods.

### Simulations to estimate the relationship between a number of tested cells and DE detection

To assess how DE detection using quasi-likelihood testing from `edgeR` depends on the number of cells, we used the R package `splatter` that simulates scRNA-seq counts with the desired properties (Zappia, Phipson, and Oshlack 2017). To estimate parameters for the simulations, we used WT mouse embryo data from a single sample of developmental stage E8.5. Additionally, we restricted the analysis to 4000 highly variable genes. We then simulated the main dataset, which contained two cell groups (imitating control and case samples, the probability to be assigned in each group is 50%) in which we assigned 25% of genes to be DE (meanlog of the effect size was assigned to 1). Additionally, we required both cell groups to have 10 batches (i.e. replicas). To estimate how the number of replicas and the imbalance of the number of replicas between control and case groups affects the detection, we then subsampled 9 datasets from the main dataset, by randomly selecting either 2, 6, or 10 samples from both control and case samples. For each subsampled dataset, we then subsampled a random number of cells (higher than 50 and lower than min(3000, number of cells in the dataset)) and estimated DE detection for the selected cells (all genes that had p-value < 0.05 were assigned as positives). For each of the 9 subsampled datasets, we downsampled cells 2000 times, and to estimate the overall trend between the number of cells and DE detection parameters (sensitivity, specificity, FDR, and False Omission Rate), we calculated the running median across 2000 downsamplings. Finally, to assess how DE detection scales with the effect size, we performed a similar procedure, but when varying the meanlog parameter in c(0.5, 1, 2, 3).

### Estimation of the relationship between the assignment (i.e. *order-k*) and neighbourhood size distribution

To estimate how neighbourhood size distribution depends on the neighbourhood assignment, we used WT mouse embryo data (developmental stage E8.5; using the provided MNN-corrected PCs as latent space) and for a wide range of [*order-k*] combinations performed neighbourhood assignment. Specifically, for *order* = 1, we used *k* = seq(50, 500, 50), and for *order* = 2, we used *k* = seq(5, 50, 5).

### Analysis of how different orders affect the sensitivity of DE detection

1. *Dataset.* We used mouse gastrulation scRNA-seq data from (Pijuan-Sala et al. 2019). As WT cells (control), we use all samples from the development stage E8.5. As ChimeraWT cells (case), we use tD-tomato negative cells from Tal1-chimera mouse embryos (which are also assigned with developmental stage E8.5). Accordingly, we concatenated counts from both experiments, and embeddings were calculated on log-normalised counts (in batch aware manner, using `batchelor::multiBatchNorm`). For the latent embedding, we used scArches integration.
2. *Selection of cell types.* To select rare cell types with various degrees of abundance, we used the provided cell type labels. Additionally, we further sub-clustered cells annotated as Forebrain/Midbrain/Hindbrain using Louvain clustering and based on the expression of known marker genes *Shh*, *Rax*, *Six3*, *Otx2*, *En1*, *Hoxb2*, *Hoxa2*, *Gbx2* (Davis et al. 1988; Simeone et al. 1992; Bouillet et al. 1995; Millet et al. 1999; Bayly, Brown, and Agarwala 2012; Lohoff et al. 2021) we annotated brain sub-clusters with one of the following (sub)cell types: Forebrain, Midbrain, Hindbrain, and Floor plate. We then selected 3 rare cell types spanning a range of relative abundance: PGC, Definitive endoderm, and Floor plate.
3. *Neighbourhood assignments.* We used the following range of [*order-k*] combinations to assign cells to neighbourhoods: for *order* = 1*, k* = c(75,150,200,250,350); for *order* = 2*, k* = c(10,15,20,25,30,40). For each [*order-k*] combination we performed 3 separate neighbourhood assignments.
4. *Per neighbourhood estimation of cell type purity.* For any given cell type, we calculated a per neighbourhood cell type purity score as the fraction of cells from the neighbourhood annotated with the cell type (relative to the total number of cells in the neighbourhood). For any given neighbourhood assignment, we then calculated the maximum cell type purity score (across all neighbourhoods from the assignment) as a proxy of how specifically the assignment results in grouping the cells from the cell type in question.
5. *Simulation of counts to supply ground truth DE.* To simulate differences in counts in a targeted manner, for each of the selected cell types, we first identified candidate genes as genes that are not DE between the conditions when tested within all abundant cell types (more than 50 cells in total). For each cell type, we then selected 40 genes and added a random number of counts (from 0 to 2) to cells from the control condition and the cell type. In addition, to ensure a wide range for logFC, we ensured a varying base expression for the selected genes. Finally, we estimated logFC for each gene/cell type, and focused the analysis on genes for which the absolute estimated logFC varies between 1 and 6. In total, we simulated 35 genes for PGC, 21 genes for Floor plate and 19 genes for Definitive endoderm.
6. *Analysis of how sensitivity in DE detection depends on cell type purity and the absolute number of cells from the cell type.* We applied miloDE for each neighbourhood assignment and, for each neighbourhood and ‘perturbed’ cell type, we estimated its cell type purity, the absolute number of cells from the cell type, and DE detection power (i.e. fraction of associated ‘perturbed’ genes that show p-value < 0.05).

### Analysis of the comparison between standard and refined neighbourhood assignments

To assess whether refined neighbourhood assignment minimises the number of neighbourhoods while ensuring complete coverage of cells with at least one neighbourhood, we used WT mouse embryo data (developmental stage E8.5; using the provided MNN-corrected PCs as latent space). We fixed *order* = 2, and for *k* = seq(10, 50, 10) performed neighbourhood assignments, followed by the refinement step (for each *k*, we performed the procedure 5 times). Then, for each neighbourhood assignment, we calculated the total number of neighbourhoods and performed matched (for the number of neighbourhoods) assignments without the refinement step. For each comparison (i.e. each we calculated the fraction of cells that did not get assigned to any neighbourhoods.

### Analysis of how AUC distribution from Augur-based classifiers depends on DE

To assess how the AUC distribution depends on whether DE is present between two tested groups, we used the R package `splatter` to simulate scRNA-seq counts (Zappia, Phipson, and Oshlack 2017). To estimate parameters for the simulations, we used WT mouse embryo data from a single sample of developmental stage E8.5. Additionally, we restricted the analysis to 4000 highly variable genes. We then simulated several datasets (54 in total), containing two tested conditions; we used 5 replicates for each condition:

- Fraction of DE genes = 0, 5 or 25%.
- Fraction of cells in control condition = 25, 50 or 75%.
- Effect size (de_facLoc) = 1,2.
- Without batch effect, with balanced batch effect (identical for one case-control replicate pair) and with unbalanced batch effect. For the unbalanced batch effect, we subsetted datasets with balanced batch effect by randomly discarding 2 case and 2 control replicates.

For each dataset we then randomly sampled cells 100 times (ensuring that the sample size lies between 75 and 500), and for each subsampling we calculated the AUC for the classifiers that separate two conditions.

### Analysis of the performance of miloDE on simulated data

1. *Dataset.* We utilised the mouse gastrulation scRNA-seq data from (Pijuan-Sala et al. 2019). As WT cells, we considered all samples from the E8.5 development stage. As ChimeraWT cells, we use tD-tomato negative cells from Tal1-chimera mouse embryos (which are also sampled from E8.5). We concatenated counts from both experiments, and embeddings were calculated on log-normalised counts (in a batch aware manner, using `batchelor::multiBatchNorm`). For the latent embedding, we used scArches.
2. *Simulation of counts to supply ground truth DE.* To simulate differences in counts in a targeted manner, we first selected a cell type (Floor plate) in which to perturb the counts for several genes. We identified candidate genes as genes that are not DE between the conditions (i.e. not DE within all abundant cell types (number of cells > 50)), and then selected 50 genes and added a random number of counts (from 0 to 2) to cells from the control condition. In addition, to ensure a wide range of logFCs were tested, we considered genes with a wide range of base expression levels. Finally, we randomly selected 5 genes with varying effect sizes, ranging from 1 to 4.5.
3. *Assessment of DE detection per neighbourhood assignment, cell type purity threshold, and ‘perturbed’ gene.* Within each neighbourhood assignment, we calculated the cell type purity score per neighbourhood as the fraction of cells in each neighbourhood annotated as being from the cell type of interest. Accordingly, for each cell type purity threshold, all neighbourhoods with cell type purity exceeding the designated threshold are annotated as ‘ground truth’ DE. Next, for each ‘perturbed’ gene we identified neighbourhoods as DE if (corrected across neighbourhoods) its p-value was less than 0.1. We then assessed DE detection power by estimating sensitivity and FDR.

### Analysis of the phenotypical comparison between Tal1+ and Tal1-embryonic cells

1. *Dataset.* We used mouse gastrulation scRNA-seq data from (Pijuan-Sala et al. 2019), specifically Tal1 chimera data, that can be loaded with `Tal1ChimeraData()`. We used tomato-td row to identify Tal1+ and Tal1-cells. We calculated log-normalised counts using scuttle::logNormCounts (McCarthy et al. 2017). For the latent embedding, we first selected 3000 highly variable genes for Tal1+ cells, and then we used MNN on batch-corrected PCs on the selected genes.
2. *Cell type ranking by the extent of transcriptional shifts.* To assess how different cell types are affected by the lack of Tal1, we first assigned neighbourhoods across the whole dataset (*order* = 2*, k* = 25). We then retained only variable genes (by selecting genes with positive variance based on `scran::modelGeneVar` estimates), and applied miloDE testing; within each neighbourhood we tested only for genes that were expressed in at least some cells (using `min_count = 3`). To estimate the extent of the transcriptional shift for each neighbourhood, we calculated two metrics: the number of DE genes (p-value corrected across genes < 0.1) and the number of ‘specifically’ DE genes. To calculate number of specifically DE genes, for each gene we first z-normalised corrected across the neighbourhoods p-values. Accordingly, a gene-neighbourhood combination is denoted as specifically DE, if its z-normalised p-value is below −3, and for each neighbourhood we calculated the total number of genes with a z-normalised p-value < −3. Note, that for all gene-neighbourhoods combinations that were not tested, we assigned p-value to 1. Finally, we assigned each neighbourhood the most enriched cell type label across the neighbourhood’s cells, and for each cell type, we calculated the distribution (across corresponding neighbourhoods) of the number of DE genes (total and neighbourhood-specific).
3. *miloDE analysis of cells contributing to blood lineage.* To systematically assess how the absence of Tal1 transcriptionally manifests in cells contributing to a blood lineage, we selected cells annotated as Haematoendothelial progenitors or Endothelium and assigned neighbourhoods (*order* = 2, *k* = 20). We then retained only variable genes (by selecting genes with positive variance based on `scran::modelGeneVar` estimates), and applied miloDE testing; within each neighbourhood we tested only for genes that were expressed in at least some cells (using `min_count = 3`).
4. *Per neighbourhood estimation of the proximity to blood progenitors.* For each cell annotated as Haematoendothelial progenitors or Endothelium, we calculated the minimum distance (in PC space) to cells that were annotated as Blood Progenitors. Accordingly, for each neighbourhood, we calculated the average (across its cells) of these minimum distances.
5. *Identification of co-regulated gene modules.* We adapted the WGCNA framework to identify gene modules consisting of co-regulated genes. Instead of expression vectors, we used logFC values (across neighbourhoods). Additionally, to minimise the input from the neighbourhoods that are not DE, we assigned all logFC values to 0 if the corrected across neighbourhoods p-value > 0.1. Additionally, for each gene-neighbourhood combination that was not tested, we assigned logFC values to 0 and p-values to 1. We restricted our analysis to genes that are DE in at least 2 neighbourhoods. Finally, to identify gene modules, we used the R package `scWGCNA` which is specifically tailored to handle scRNA-seq data (Feregrino and Tschopp 2022). Specifically, we employed `scWGCNA::run.scWGCNA` using neighbourhoods instead of single cells (and therefore skipping the calculation of pseudocells), otherwise with the default settings.
6. *Gene ontology enrichment analysis.* To assess which biological processes are enriched across different gene modules, we used `enrichR::enrichr` (within `GO_Biological_Process_2021` database) (E. Y. Chen et al. 2013; Kuleshov et al. 2016). For each gene module, all gene ontologies with adjusted p-value < 0.1 were assigned as significantly enriched.

### Analysis of macrophage-specific transcriptional shifts upon Idiopathic Pulmonary Fibrosis

1. *Dataset.* 4 datasets, containing cells from both healthy and IPF donors (Morse et al. 2019; Reyfman et al. 2019; Habermann et al. 2020; Adams et al. 2020) were downloaded from the original source and mapped onto the Azimuth lung reference (available at https://cellxgene.cziscience.com/e/f72958f5-7f42-4ebb-98da-445b0c6de516.cxg/). To assign neighbourhoods, we used provided latent embedding that are generated by the Azimuth integration pipeline.
2. *Neighbourhood assignment and DE testing.* To characterise macrophage-specific transcriptional shifts, we selected all cells annotated as macrophage, assigned neighbourhoods (*order* = 2, *k* = 30). We then retained only variable genes (by selecting genes with positive variance based on `scran::modelGeneVar` estimates), and applied miloDE testing, using the dataset ID as a covariate. Within each neighbourhood, we tested genes that are expressed in macrophage annotated cells (gene selection was performed based on the output of `edgeR::filterByExpr`, `min_count = 3`). Additionally, within each neighbourhood we discarded donors, for whom we had less than 3 cells in the neighbourhood.
3. *Identification of gene sets.* To identify macrophage-specific gene sets, we clustered genes using Louvain clustering. We restricted the analysis to genes that were strongly upregulated in IPF donors. Specifically, we selected genes that satisfied the next criteria:

- Corrected across neighbourhoods p-value < 0.05 in at least 10% of the neighbourhoods.
- Absolute average logFC across significant (p-value (corrected across neighbourhoods) < 0.05) neighbourhoods > 1.
- Gene is up-regulated in IPF donors (i.e. has negative logFC) in at least 75% of neighbourhoods. As a vector for each gene, we use logFC (across neighbourhoods), with logFC being set to 0 if the corrected across the neighbourhoods p-value > 0.05 (henceforth referred to as corrected logFC). We then computed the shared nearest neighbours graph (on the first 5 PCs, using `scran::buildSNNGraph`) and Louvain clustering was calculated using `igraph::cluster_louvain` (Csardi, Nepusz, and Others 2006; Blondel et al. 2008) (resolution = 1).
4. *Per gene set discovery of ‘marker’ neighbourhoods.* To identify the neighbourhoods in which genes between different gene sets are DE, we used `scran::findMarkers` on logFC vectors (across neighbourhoods; using corrected logFC). We then selected the ‘top’ 3 neighbourhoods per gene set, where top neighbourhoods were defined by the ‘Top’ column in `scran::findMarkers` output and represent a minimal number of neighbourhoods required to separate any cluster from any other cluster (specified by `pval.type = “any”`).
5. *Characterisation of DE patterns based on the prevalence of DE neighbourhoods in different stages of fibrotic progression.* To characterise whether a gene is DE preferably in the neighbourhoods that are significantly enriched for IPF cells or in the neighbourhoods that contain a comparable number of cells from healthy and IPF donors, we restricted the analysis to neighbourhoods with negative logFC for DA test and split them into two groups: neighbourhoods that are significantly enriched for IPF cells (SpatialFDR for DA test < 0.05) and neighbourhoods that contain a comparable number of cells from healthy and IPF donors (SpatialFDR for DA test >= 0.05). Then, for each gene, we extracted the number of DE neighbourhoods (p-value corrected across neighbourhoods < 0.05) in each of the groups, and performed a Fisher test to assess whether any of the two neighbourhood groups contain significantly more DE neighbourhoods. We performed a Fisher test for all genes that we used for the Louvain clustering and used a corrected p-value (cutoff of 0.05) to decide whether the difference is significant or not. Accordingly, we split all genes into 3 classes: genes that are DE significantly more frequently in either of the two groups and genes that are DE relatively equally in both groups.
6. *Comparison with per cell type DE estimation.* To assess whether genes that we identify with miloDE as upregulated in IPF are also DE on a whole cell type level, we implemented the same edgeR framework but across all macrophages (we performed testing only for genes that we used for the clustering i.e. genes that are strongly upregulated in IPF donors). We then assigned genes with p-value < 0.05 as significantly DE on the cell type level.
7. *Per gene set characterisation of average DE pattern along the fibrosis progression.* To characterise per gene set DE patterns along the fibrosis progression, we grouped neighbourhoods based on their logFC for the DA test into 50 equal-sized bins. For each set and logFC-DA bin, we then calculated the average (across all aggregated neighbourhoods and genes from the gene set) corrected logFC across all genes from the set and average (across all aggregated neighbourhoods) fraction of genes (from the gene set) that are significantly DE in the corresponding neighbourhood.
8. *Gene ontology enrichment analysis.* To assess which biological processes are enriched across different gene modules, we used `enrichR::enrichr` (within `GO_Biological_Process_2021` database). For each gene set, we then selected the top 5 gene ontologies (based on adjusted p-value) and for the union of all top gene ontologies, we estimated how significantly each of them is enriched in each gene set.

## Acknowledgments

We thank Mike Morgan and Alice Kluzer for optimising and scaling the original Milo approach which enabled a scalable implementation of miloDE. We also thank Jean Fan, Brendan Miller, Austin Hartman, and Gesmira Molla for fruitful discussions that helped to shape the method and interpretation of the results. We thank Berthold Göttgens, Luke Harland, and Bart Theeuwes for the discussions and valuable feedback on the implementation of miloDE in the context of chimeric mouse embryos lacking Tal1. We thank Ana-Maria Cujba and Amanda Oliver for the discussions and valuable feedback on the implementation of miloDE in the context of macrophage activation in IPF.

## Author contribution

A.M., R.S. and J.M. conceived the method. A.M. developed the method, wrote the code, and performed the analysis, with input from E.D. and L.R.. All authors interpreted the results. A.M. and J.M. wrote the manuscript, with input from all the authors. J.M. and R.S. oversaw the project.

## Ethics approval and consent to participate

Not applicable.

## Consent for publication

Not applicable.

## Additional files

Additional file 1: Table S1. List of enriched gene ontologies for each module detected in Tal1-analysis.

Additional file 2: Supplementary Notes.

## Spplementary figures

**Supplementary Figure 1.**
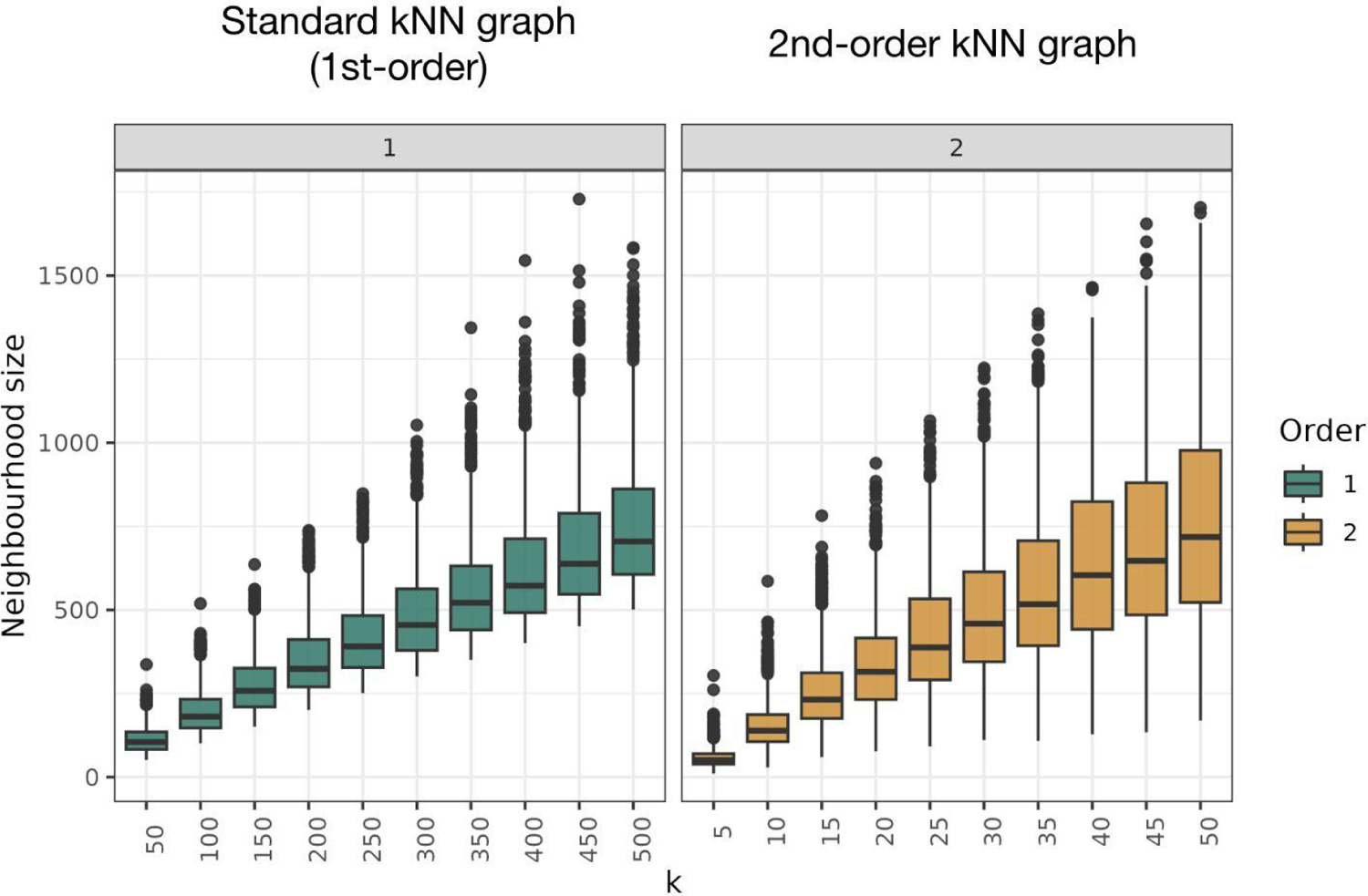
The estimate of the scaling between *k* (x-axis) and the neighbourhood size distribution (y-axis) for either 1st- or 2nd-order KNN graphs (in facets and in colour)

**Supplementary Figure 2.**
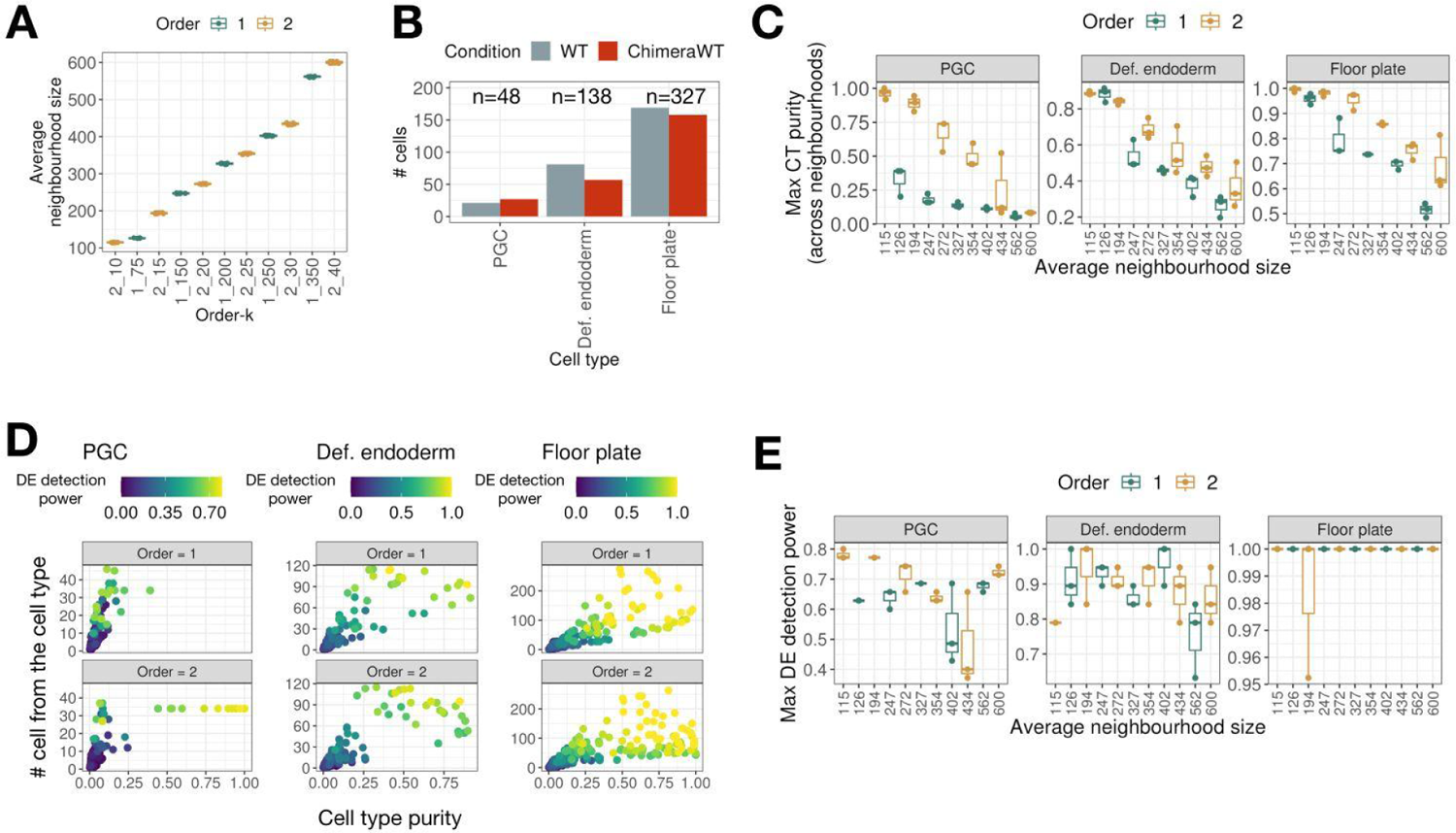
2nd-order neighbourhood graphs preserve neighbourhood homogeneity better while also controlling for the average neighbourhood size. A. Boxplot representing the relationship between average neighbourhood size and the selected *order-k* grid that we used to perform neighbourhoods assignments. B. Barplot representing cell types composition across cell types and conditions. Total number of cells per cell type is indicated above each cell type. C. Boxplots representing maximum cell type purity (across neighbourhoods) for each cell type and each neighbourhood assignment. x-axis correspond to the average neighbourhood size, colours correspond to the order of the graph. D. Scater plot representing the relationship between cell type purity (x-axis), number of cells from the cell type (y-axis) and DE detection power (in colour). Each point corresponds to one neighbourhood. E. Boxplot representing the relationship between an assignment (i.e. *order-k*) and maximum DE detection power (across neighbourhoods).

**Supplementary Figure 3.**
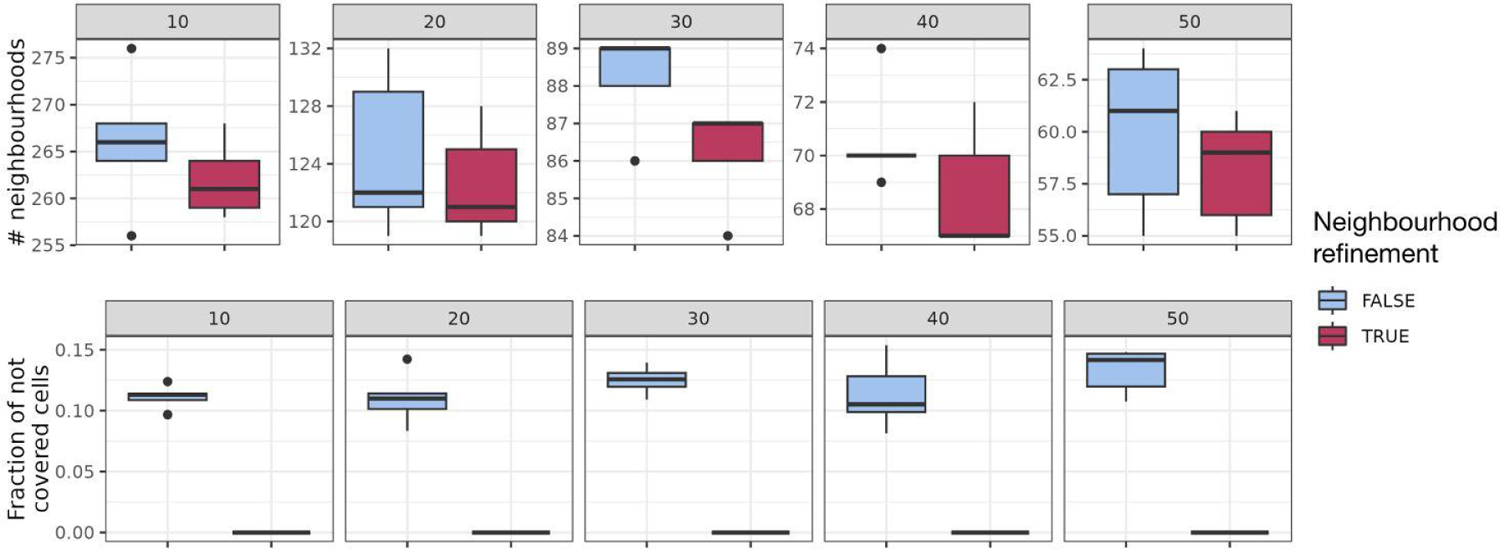
Post hoc neighbourhood filtering minimises the number of assigned neighbourhoods while also controlling for the inclusion of all cells in at least one neighbourhood. Each facet corresponds to a different *k* in the graph assignment (using 2nd-order graph), and colours correspond to whether filtering was performed or not. Top panels correspond to the number of neighbourhoods that is matched between two filtering options, and bottom panels correspond to the fraction of cells that are not assigned with any neighbourhoods.

**Supplementary Figure 4.**
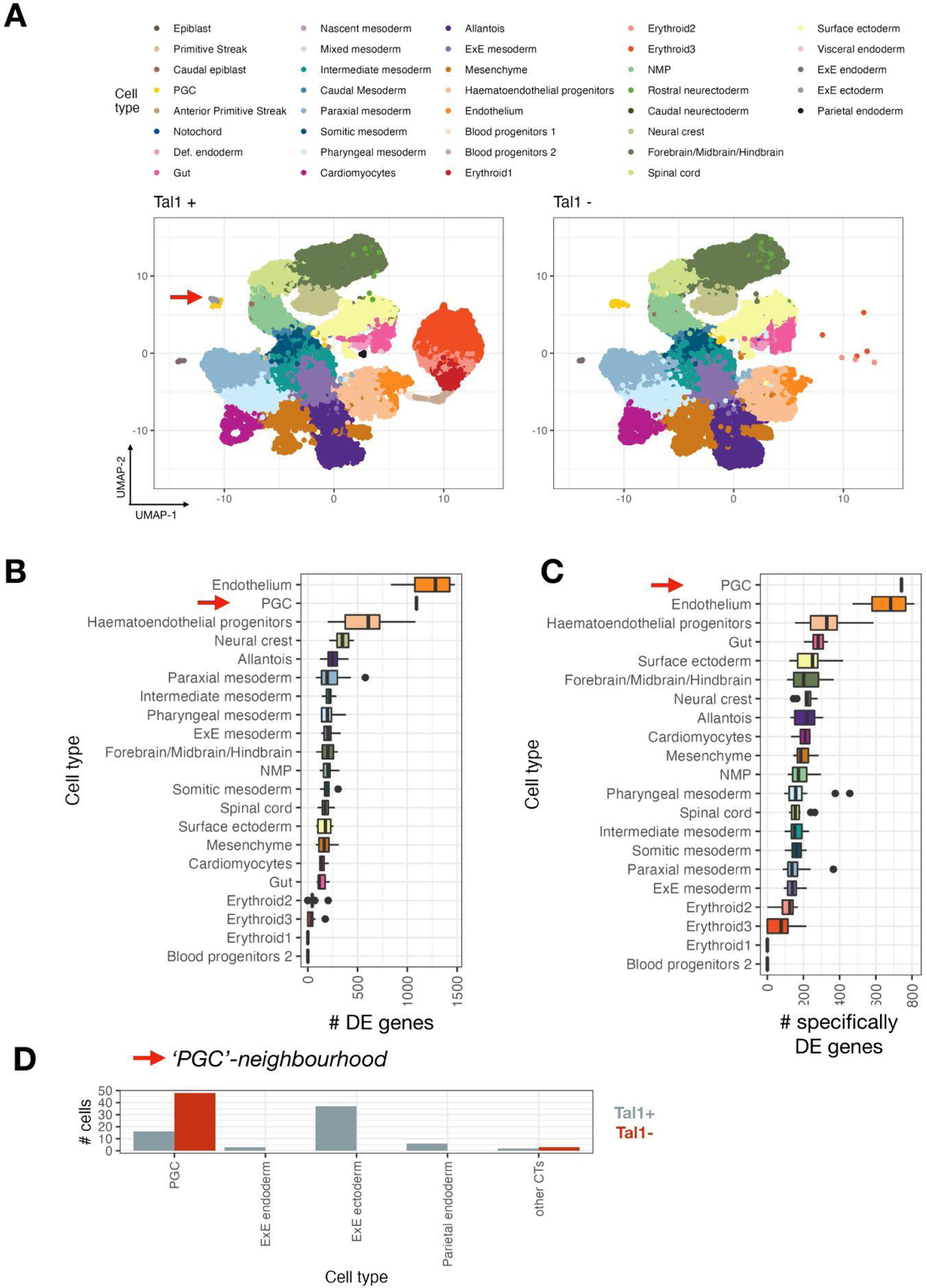
Cells contributing to blood lineage have higher ‘degree of perturbation’. A. UMAP representing a manifold of chimera mouse embryos, colours correspond to the cell types, and facets correspond to whether cells carry a knockout of Tal1. B. Boxplots representing cell type ranking according to the number of DE genes per neighbourhood, with neighbourhoods being grouped by associated cell type. C. Boxplots representing cell type ranking according to the number of specifically DE genes (with respect to other neighbourhoods) per neighbourhood, with neighbourhoods being grouped by associated cell type. D. Barplot representing breakdown by cell types and conditions for the PGC-neighbourhood that shows a high ‘degree of perturbation’.

**Supplementary Figure 5.**
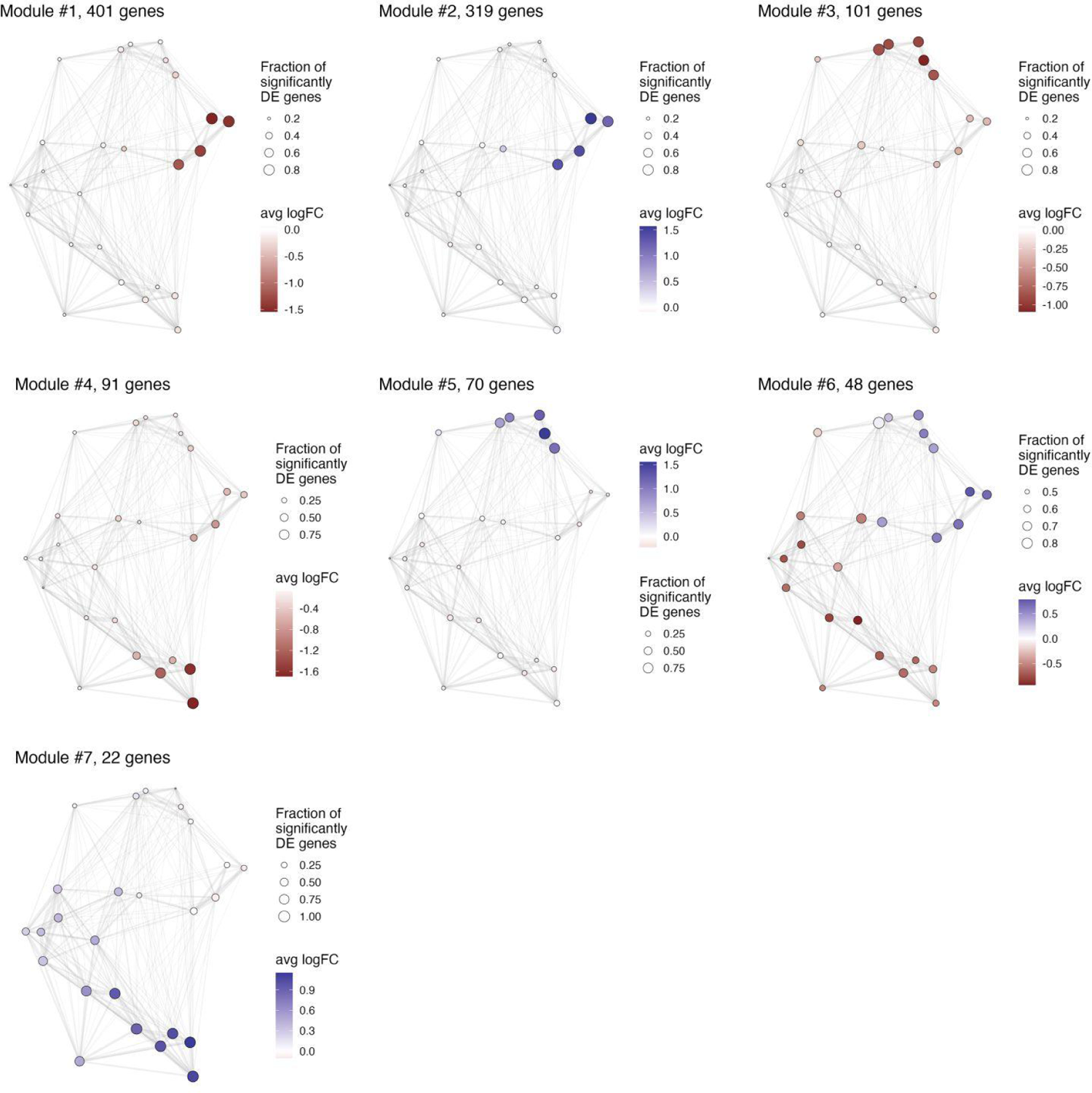
‘Neighbourhood’ graphs representing transcriptional profiles for different DE patterns in chimeric mouse embryos lacking Tal1.

**Supplementary Figure 6.**
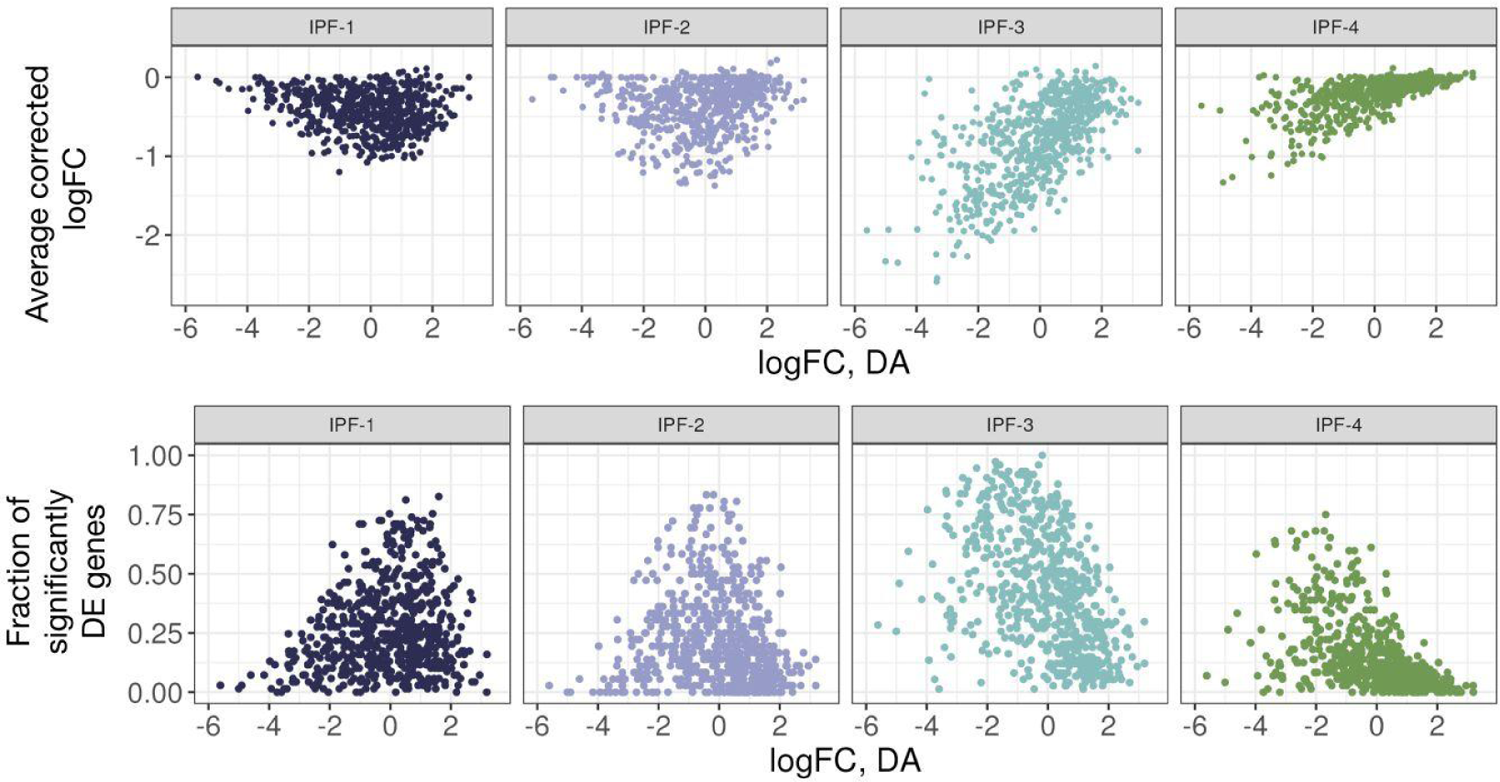
Gene sets vary in their ‘location’ and effect size distributions. Scatter plots represent the relationship between logFC, DA (i.e. proxy for phenotypic fibrosis state, x-axis) and average corrected logFC (upper panels) and fraction of significantly DE genes (lower panels). Each point corresponds to one neighbourhood, facets correspond to different gene sets.

## References

Adams, Taylor S., Jonas C. Schupp, Sergio Poli, Ehab A. Ayaub, Nir Neumark, Farida Ahangari, Sarah G. Chu, et al. 2020. “Single-Cell RNA-Seq Reveals Ectopic and Aberrant Lung-Resident Cell Populations in Idiopathic Pulmonary Fibrosis.” Science Advances 6 (28): eaba1983.

Alber, Andreas, Sarah E. M. Howie, William A. H. Wallace, and Nikhil Hirani. 2012. “The Role of Macrophages in Healing the Wounded Lung.” International Journal of Experimental Pathology 93 (4): 243–51.

Amezquita, Robert A., Aaron T. L. Lun, Etienne Becht, Vince J. Carey, Lindsay N. Carpp, Ludwig Geistlinger, Federico Marini, et al. 2020. “Orchestrating Single-Cell Analysis with Bioconductor.” Nature Methods 17 (2): 137–45.

Baran, Yael, Akhiad Bercovich, Arnau Sebe-Pedros, Yaniv Lubling, Amir Giladi, Elad Chomsky, Zohar Meir, Michael Hoichman, Aviezer Lifshitz, and Amos Tanay. 2019. “MetaCell: Analysis of Single-Cell RNA-Seq Data Using K-Nn Graph Partitions.” Genome Biology 20 (1): 206.

Bayly, Roy D., Charmaine Y. Brown, and Seema Agarwala. 2012. “A Novel Role for FOXA2 and SHH in Organizing Midbrain Signaling Centers.” Developmental Biology 369 (1): 32–42.

Blondel, Vincent D., Jean-Loup Guillaume, Renaud Lambiotte, and Etienne Lefebvre. 2008. “Fast Unfolding of Communities in Large Networks.” Journal of Statistical Mechanics 2008 (10): P10008.

Borthwick, L. A. 2016. “The IL-1 Cytokine Family and Its Role in Inflammation and Fibrosis in the Lung.” Seminars in Immunopathology 38 (4): 517–34.

Bouillet, P., C. Chazaud, M. Oulad-Abdelghani, P. Dollé, and P. Chambon. 1995. “Sequence and Expression Pattern of the Stra7 (Gbx-2) Homeobox-Containing Gene Induced by Retinoic Acid in P19 Embryonal Carcinoma Cells.” Developmental Dynamics: An Official Publication of the American Association of Anatomists 204 (4): 372–82.

Bourgon, Richard, Robert Gentleman, and Wolfgang Huber. 2010. “Independent Filtering Increases Detection Power for High-Throughput Experiments.” Proceedings of the National Academy of Sciences of the United States of America 107 (21): 9546–51.

Boyeau, Pierre, Jeffrey Regier, Adam Gayoso, Michael I. Jordan, Romain Lopez, and Nir Yosef. n.d. “An Empirical Bayes Method for Differential Expression Analysis of Single Cells with Deep Generative Models.” https://doi.org/10.1101/2022.05.27.493625.

Chagraoui, Hedia, Maiken S. Kristiansen, Juan Pablo Ruiz, Ana Serra-Barros, Johanna Richter, Elisa Hall-Ponselé, Nicki Gray, et al. 2018. “SCL/TAL1 Cooperates with Polycomb RYBP-PRC1 to Suppress Alternative Lineages in Blood-Fated Cells.” Nature Communications 9 (1): 5375.

Chen, Edward Y., Christopher M. Tan, Yan Kou, Qiaonan Duan, Zichen Wang, Gabriela Vaz Meirelles, Neil R. Clark, and Avi Ma’ayan. 2013. “Enrichr: Interactive and Collaborative HTML5 Gene List Enrichment Analysis Tool.” BMC Bioinformatics 14 (April): 128.

Chen, Yunshun, Aaron T. L. Lun, and Gordon K. Smyth. 2016. “From Reads to Genes to Pathways: Differential Expression Analysis of RNA-Seq Experiments Using Rsubread and the edgeR Quasi-Likelihood Pipeline.” F1000Research. https://doi.org/10.12688/f1000research.8987.1.

Crowell, Helena L., Charlotte Soneson, Pierre-Luc Germain, Daniela Calini, Ludovic Collin, Catarina Raposo, Dheeraj Malhotra, and Mark D. Robinson. 2020. “Muscat Detects Subpopulation-Specific State Transitions from Multi-Sample Multi-Condition Single-Cell Transcriptomics Data.” Nature Communications 11 (1): 6077.

Csardi, Gabor, Tamas Nepusz, and Others. 2006. “The Igraph Software Package for Complex Network Research.” InterJournal, Complex Systems 1695 (5): 1–9.

Dann, Emma, Neil C. Henderson, Sarah A. Teichmann, Michael D. Morgan, and John C. Marioni. 2021. “Differential Abundance Testing on Single-Cell Data Using K-Nearest Neighbor Graphs.” Nature Biotechnology, September. https://doi.org/10.1038/s41587-021-01033-z.

Dann, Emma, Sarah A. Teichmann, and John C. Marioni. 2022. “Precise Identification of Cell States Altered in Disease with Healthy Single-Cell References.” bioRxiv. https://doi.org/10.1101/2022.11.10.515939.

Davis, C. A., S. E. Noble-Topham, J. Rossant, and A. L. Joyner. 1988. “Expression of the Homeo Box-Containing Gene En-2 Delineates a Specific Region of the Developing Mouse Brain.” Genes & Development 2 (3): 361–71.

Delmans, Mihails, and Martin Hemberg. 2016. “Discrete Distributional Differential Expression (D3E)--a Tool for Gene Expression Analysis of Single-Cell RNA-Seq Data.” BMC Bioinformatics 17 (February): 110.

Dumon, Stephanie, David S. Walton, Giacomo Volpe, Nicola Wilson, Emilie Dassé, Walter Del Pozzo, Josette-Renee Landry, et al. 2012. “Itga2b Regulation at the Onset of Definitive Hematopoiesis and Commitment to Differentiation.” PloS One 7 (8): e43300.

Eddelbuettel, Dirk. n.d. Seamless R and C++ Integration with Rcpp. Springer New York. Accessed January 20, 2023.

Eddelbuettel, Dirk, and Romain Francois. 2011. “Rcpp: Seamless R and C++ Integration.” Journal of Statistical Software 40 (April): 1–18.

Elmentaite, Rasa, Alexander D. B. Ross, Kenny Roberts, Kylie R. James, Daniel Ortmann, Tomás Gomes, Komal Nayak, et al. 2020. “Single-Cell Sequencing of Developing Human Gut Reveals Transcriptional Links to Childhood Crohn’s Disease.” Developmental Cell 55 (6): 771–83.e5.

Fastrès, Aline, Florence Felice, Elodie Roels, Catherine Moermans, Jean-Louis Corhay, Fabrice Bureau, Renaud Louis, Cécile Clercx, and Julien Guiot. 2017. “The Lung Microbiome in Idiopathic Pulmonary Fibrosis: A Promising Approach for Targeted Therapies.” International Journal of Molecular Sciences 18 (12). https://doi.org/10.3390/ijms18122735.

Feregrino, Christian, and Patrick Tschopp. 2022. “Assessing Evolutionary and Developmental Transcriptome Dynamics in Homologous Cell Types.” Developmental Dynamics: An Official Publication of the American Association of Anatomists 251 (9): 1472–89.

Finak, Greg, Andrew McDavid, Masanao Yajima, Jingyuan Deng, Vivian Gersuk, Alex K. Shalek, Chloe K. Slichter, et al. 2015. “MAST: A Flexible Statistical Framework for Assessing Transcriptional Changes and Characterizing Heterogeneity in Single-Cell RNA Sequencing Data.” Genome Biology 16 (December): 278.

Gagnon, Jake, Lira Pi, Matthew Ryals, Qingwen Wan, Wenxing Hu, Zhengyu Ouyang, Baohong Zhang, and Kejie Li. 2022. “Recommendations of scRNA-Seq Differential Gene Expression Analysis Based on Comprehensive Benchmarking.” Life 12 (6). https://doi.org/10.3390/life12060850.

Griffiths J, Lun A (2022). *MouseGastrulationData:* Single-Cell -omics Data across Mouse Gastrulation and Early Organogenesis. R package version 1.12.0, https://github.com/MarioniLab/MouseGastrulationData.

Gut, Gabriele, Michelle D. Tadmor, Dana Pe’er, Lucas Pelkmans, and Prisca Liberali. 2015. “Trajectories of Cell-Cycle Progression from Fixed Cell Populations.” Nature Methods 12 (10): 951–54.

Habermann, Arun C., Austin J. Gutierrez, Linh T. Bui, Stephanie L. Yahn, Nichelle I. Winters, Carla L. Calvi, Lance Peter, et al. 2020. “Single-Cell RNA Sequencing Reveals Profibrotic Roles of Distinct Epithelial and Mesenchymal Lineages in Pulmonary Fibrosis.” Science Advances 6 (28): eaba1972.

Haghverdi, Laleh, Aaron T. L. Lun, Michael D. Morgan, and John C. Marioni. 2018. “Batch Effects in Single-Cell RNA-Sequencing Data Are Corrected by Matching Mutual Nearest Neighbors.” Nature Biotechnology 36 (5): 421–27.

Hao, Yuhan, Stephanie Hao, Erica Andersen-Nissen, William M. Mauck 3rd, Shiwei Zheng, Andrew Butler, Maddie J. Lee, et al. 2021. “Integrated Analysis of Multimodal Single-Cell Data.” Cell 184 (13): 3573–87.e29.

Hurk, Mark van den, Shong Lau, Maria C. Marchetto, Jerome Mertens, Shani Stern, Olga Corti, Alexis Brice, et al. 2022. “Druggable Transcriptomic Pathways Revealed in Parkinson’s Patient-Derived Midbrain Neurons.” Npj Parkinson’s Disease. https://doi.org/10.1038/s41531-022-00400-0.

Kharchenko, Peter V., Lev Silberstein, and David T. Scadden. 2014. “Bayesian Approach to Single-Cell Differential Expression Analysis.” Nature Methods 11 (7): 740–42.

Korthauer, Keegan D., Li-Fang Chu, Michael A. Newton, Yuan Li, James Thomson, Ron Stewart, and Christina Kendziorski. 2016. “A Statistical Approach for Identifying Differential Distributions in Single-Cell RNA-Seq Experiments.” Genome Biology. https://doi.org/10.1186/s13059-016-1077-y.

Kuleshov, Maxim V., Matthew R. Jones, Andrew D. Rouillard, Nicolas F. Fernandez, Qiaonan Duan, Zichen Wang, Simon Koplev, et al. 2016. “Enrichr: A Comprehensive Gene Set Enrichment Analysis Web Server 2016 Update.” Nucleic Acids Research 44 (W1): W90–97.

Langfelder, Peter, and Steve Horvath. 2008. “WGCNA: An R Package for Weighted Correlation Network Analysis.” BMC Bioinformatics 9 (December): 559.

Langfelder, Peter, Rui Luo, Michael C. Oldham, and Steve Horvath. 2011. “Is My Network Module Preserved and Reproducible?” PLoS Computational Biology. https://doi.org/10.1371/journal.pcbi.1001057.

Lederer, David J., and Fernando J. Martinez. 2018. “Idiopathic Pulmonary Fibrosis.” The New England Journal of Medicine.

Levine, Jacob H., Erin F. Simonds, Sean C. Bendall, Kara L. Davis, El-Ad D. Amir, Michelle D. Tadmor, Oren Litvin, et al. 2015. “Data-Driven Phenotypic Dissection of AML Reveals Progenitor-like Cells That Correlate with Prognosis.” Cell 162 (1): 184–97.

Lohoff, T., S. Ghazanfar, A. Missarova, N. Koulena, N. Pierson, J. A. Griffiths, E. S. Bardot, et al. 2021. “Integration of Spatial and Single-Cell Transcriptomic Data Elucidates Mouse Organogenesis.” Nature Biotechnology, September. https://doi.org/10.1038/s41587-021-01006-2.

Lotfollahi, Mohammad, Mohsen Naghipourfar, Malte D. Luecken, Matin Khajavi, Maren Büttner, Marco Wagenstetter, Žiga Avsec, et al. 2022. “Mapping Single-Cell Data to Reference Atlases by Transfer Learning.” Nature Biotechnology 40 (1): 121–30.

Love, Michael I., Wolfgang Huber, and Simon Anders. 2014. “Moderated Estimation of Fold Change and Dispersion for RNA-Seq Data with DESeq2.” Genome Biology 15 (12): 550.

Lun, Aaron T. L., Yunshun Chen, and Gordon K. Smyth. 2016. “It’s DE-Licious: A Recipe for Differential Expression Analyses of RNA-Seq Experiments Using Quasi-Likelihood Methods in edgeR.” Methods in Molecular Biology 1418: 391–416.

Lun, Aaron T. L., and John C. Marioni. 2017. “Overcoming Confounding Plate Effects in Differential Expression Analyses of Single-Cell RNA-Seq Data.” Biostatistics 18 (3): 451–64.

Lun, Aaron T. L., Davis J. McCarthy, and John C. Marioni. 2016. “A Step-by-Step Workflow for Low-Level Analysis of Single-Cell RNA-Seq Data with Bioconductor.” F1000Research. https://doi.org/10.12688/f1000research.9501.2.

Lun, Aaron T. L., Arianne C. Richard, and John C. Marioni. 2017. “Testing for Differential Abundance in Mass Cytometry Data.” Nature Methods 14 (7): 707–9.

Madala, Satish K., Stephanie Schmidt, Cynthia Davidson, Machiko Ikegami, Susan Wert, and William D. Hardie. 2012. “MEK-ERK Pathway Modulation Ameliorates Pulmonary Fibrosis Associated with Epidermal Growth Factor Receptor Activation.” American Journal of Respiratory Cell and Molecular Biology 46 (3): 380–88.

McCarthy, Davis J., Kieran R. Campbell, Aaron T. L. Lun, and Quin F. Wills. 2017. “Scater: Pre-Processing, Quality Control, Normalization and Visualization of Single-Cell RNA-Seq Data in R.” Bioinformatics 33 (8): 1179–86.

McCarthy, Davis J., Yunshun Chen, and Gordon K. Smyth. 2012. “Differential Expression Analysis of Multifactor RNA-Seq Experiments with Respect to Biological Variation.” Nucleic Acids Research 40 (10): 4288–97.

McGrath, Kathleen E., Jenna M. Frame, Katherine H. Fegan, James R. Bowen, Simon J. Conway, Seana C. Catherman, Paul D. Kingsley, Anne D. Koniski, and James Palis. 2015. “Distinct Sources of Hematopoietic Progenitors Emerge before HSCs and Provide Functional Blood Cells in the Mammalian Embryo.” Cell Reports 11 (12): 1892–1904.

Millet, S., K. Campbell, D. J. Epstein, K. Losos, E. Harris, and A. L. Joyner. 1999. “A Role for Gbx2 in Repression of Otx2 and Positioning the Mid/hindbrain Organizer.” Nature 401 (6749): 161–64.

Montoro, Daniel T., Adam L. Haber, Moshe Biton, Vladimir Vinarsky, Brian Lin, Susan E. Birket, Feng Yuan, et al. 2018. “A Revised Airway Epithelial Hierarchy Includes CFTR-Expressing Ionocytes.” Nature. https://doi.org/10.1038/s41586-018-0393-7.

Morse, Christina, Tracy Tabib, John Sembrat, Kristina L. Buschur, Humberto Trejo Bittar, Eleanor Valenzi, Yale Jiang, et al. 2019. “Proliferating SPP1/MERTK-Expressing Macrophages in Idiopathic Pulmonary Fibrosis.” The European Respiratory Journal: Official Journal of the European Society for Clinical Respiratory Physiology 54 (2). https://doi.org/10.1183/13993003.02441-2018.

Murray, Peter J., and Thomas A. Wynn. 2011. “Protective and Pathogenic Functions of Macrophage Subsets.” Nature Reviews. Immunology 11 (11): 723–37.

Ogawa, Tatsuro, Shigeyuki Shichino, Satoshi Ueha, and Kouji Matsushima. 2021. “Macrophages in Lung Fibrosis.” International Immunology 33 (12): 665–71.

Org, Tõnis, Dan Duan, Roberto Ferrari, Amelie Montel-Hagen, Ben Van Handel, Marc A. Kerényi, Rajkumar Sasidharan, et al. 2015. “Scl Binds to Primed Enhancers in Mesoderm to Regulate Hematopoietic and Cardiac Fate Divergence.” The EMBO Journal 34 (6): 759–77.

Ottersbach, Katrin. 2019. “Endothelial-to-Haematopoietic Transition: An Update on the Process of Making Blood.” Biochemical Society Transactions 47 (2): 591–601.

Persad, Sitara, Zi-Ning Choo, Christine Dien, Ignas Masilionis, Ronan Chaligné, Tal Nawy, Chrysothemis C. Brown, Itsik Pe’er, Manu Setty, and Dana Pe’er. 2022. “SEACells: Inference of Transcriptional and Epigenomic Cellular States from Single-Cell Genomics Data.” bioRxiv. https://doi.org/10.1101/2022.04.02.486748.

Petukhov, Viktor, Anna Igolkina, Rasmus Rydbirk, Shenglin Mei, Lars Christoffersen, Konstantin Khodosevich, and Peter V. Kharchenko. 2022. “Case-Control Analysis of Single-Cell RNA-Seq Studies.” bioRxiv. https://doi.org/10.1101/2022.03.15.484475.

Pijuan-Sala, Blanca, Jonathan A. Griffiths, Carolina Guibentif, Tom W. Hiscock, Wajid Jawaid, Fernando J. Calero-Nieto, Carla Mulas, et al. 2019. “A Single-Cell Molecular Map of Mouse Gastrulation and Early Organogenesis.” Nature 566 (7745): 490–95.

Reyfman, Paul A., James M. Walter, Nikita Joshi, Kishore R. Anekalla, Alexandra C. McQuattie-Pimentel, Stephen Chiu, Ramiro Fernandez, et al. 2019. “Single-Cell Transcriptomic Analysis of Human Lung Provides Insights into the Pathobiology of Pulmonary Fibrosis.” American Journal of Respiratory and Critical Care Medicine 199 (12): 1517–36.

Ritchie, Matthew E., Belinda Phipson, Di Wu, Yifang Hu, Charity W. Law, Wei Shi, and Gordon K. Smyth. 2015. “Limma Powers Differential Expression Analyses for RNA-Sequencing and Microarray Studies.” Nucleic Acids Research 43 (7): e47.

Robb, L., N. J. Elwood, A. G. Elefanty, F. Köntgen, R. Li, L. D. Barnett, and C. G. Begley. 1996. “The Scl Gene Product Is Required for the Generation of All Hematopoietic Lineages in the Adult Mouse.” The EMBO Journal 15 (16): 4123–29.

Robinson, Mark D., Davis J. McCarthy, and Gordon K. Smyth. 2010. “edgeR: A Bioconductor Package for Differential Expression Analysis of Digital Gene Expression Data.” Bioinformatics 26 (1): 139–40.

Rood, Jennifer E., Aidan Maartens, Anna Hupalowska, Sarah A. Teichmann, and Aviv Regev. 2022. “Impact of the Human Cell Atlas on Medicine.” *Nature Medicine*, December. https://doi.org/10.1038/s41591-022-02104-7.

Scialdone, Antonio, Yosuke Tanaka, Wajid Jawaid, Victoria Moignard, Nicola K. Wilson, Iain C. Macaulay, John C. Marioni, and Berthold Göttgens. 2016. “Resolving Early Mesoderm Diversification through Single-Cell Expression Profiling.” Nature 535 (7611): 289–93.

Shivdasani, R. A., E. L. Mayer, and S. H. Orkin. 1995. “Absence of Blood Formation in Mice Lacking the T-Cell Leukaemia Oncoprotein Tal-1/SCL.” Nature 373 (6513): 432–34.

Simeone, A., D. Acampora, M. Gulisano, A. Stornaiuolo, and E. Boncinelli. 1992. “Nested Expression Domains of Four Homeobox Genes in Developing Rostral Brain.” Nature 358 (6388): 687–90.

Skinnider, Michael A., Jordan W. Squair, Claudia Kathe, Mark A. Anderson, Matthieu Gautier, Kaya J. E. Matson, Marco Milano, et al. 2021. “Cell Type Prioritization in Single-Cell Data.” Nature Biotechnology 39 (1): 30–34.

Soneson, Charlotte, and Mark D. Robinson. 2018. “Bias, Robustness and Scalability in Single-Cell Differential Expression Analysis.” Nature Methods 15 (4): 255–61.

Squair, Jordan W., Matthieu Gautier, Claudia Kathe, Mark A. Anderson, Nicholas D. James, Thomas H. Hutson, Rémi Hudelle, et al. 2021. “Confronting False Discoveries in Single-Cell Differential Expression.” Nature Communications 12 (1): 5692.

Stephenson, Emily, Gary Reynolds, Rachel A. Botting, Fernando J. Calero-Nieto, Michael D. Morgan, Zewen Kelvin Tuong, Karsten Bach, et al. 2021. “Single-Cell Multi-Omics Analysis of the Immune Response in COVID-19.” Nature Medicine 27 (5): 904–16.

Tirosh, Itay, Benjamin Izar, Sanjay M. Prakadan, Marc H. Wadsworth 2nd, Daniel Treacy, John J. Trombetta, Asaf Rotem, et al. 2016. “Dissecting the Multicellular Ecosystem of Metastatic Melanoma by Single-Cell RNA-Seq.” Science 352 (6282): 189–96.

Wang, Tianyu, Boyang Li, Craig E. Nelson, and Sheida Nabavi. 2019. “Comparative Analysis of Differential Gene Expression Analysis Tools for Single-Cell RNA Sequencing Data.” BMC Bioinformatics 20 (1): 40.

Wang, Xin-Sheng, Zachary Simmons, Wenlei Liu, Philip J. Boyer, and James R. Connor. 2006. “Differential Expression of Genes in Amyotrophic Lateral Sclerosis Revealed by Profiling the Post Mortem Cortex.” Amyotrophic Lateral Sclerosis: Official Publication of the World Federation of Neurology Research Group on Motor Neuron Diseases 7 (4): 201–10.

Yang, Ivana V., Tasha E. Fingerlin, Christopher M. Evans, Marvin I. Schwarz, and David A. Schwartz. 2015. “MUC5B and Idiopathic Pulmonary Fibrosis.” Annals of the American Thoracic Society 12 Suppl 2 (Suppl 2): S193–99.

Ye, Chengzhong, Terence P. Speed, and Agus Salim. 2019. “DECENT: Differential Expression with Capture Efficiency adjustmeNT for Single-Cell RNA-Seq Data.” Bioinformatics 35 (24): 5155–62.

Zappia, Luke, Belinda Phipson, and Alicia Oshlack. 2017. “Splatter: Simulation of Single-Cell RNA Sequencing Data.” Genome Biology 18 (1): 174.

Zhang, Lei, Yi Wang, Guorao Wu, Weining Xiong, Weikuan Gu, and Cong-Yi Wang. 2018. “Macrophages: Friend or Foe in Idiopathic Pulmonary Fibrosis?” Respiratory Research 19 (1): 170.

