## Supplementary Notes for "Sensitive cluster-free differential expression testing"

**Supplementary Note 1. Supervised embedding enables more sensitive detection of DE genes.**

Given the range of integration/embedding techniques (Chen et al. 2019; Luecken et al. 2022), it is important to consider how they might impact the ability to detect DE genes. In particular, such techniques aim to group cells that are transcriptionally similar, while DE approaches aim to identify genes that are transcriptionally distinct. This seemingly self-contradictory observation can be at least partially resolved by specifying that the task of miloDE is to identify case-specific cell states (defined as a change in expression of certain gene(s) in the given region(s) of the manifold) that are not present in the control data. Within this paradigm, we can decompose the contribution of each gene to the final manifold into a control-specific component (how much this gene contributes to the formation of the cell types in both healthy and case settings) and a case-specific component (manifestation of the gene's relevance only in the case setting). Therefore, to identify a per gene case-specific component and determine whether it is significant, the ideal embedding minimises the case-specific signal and 'maps' cells using only control-specific signatures. Driven by this, we hypothesised that supervised approaches, where models are learned on the reference (i.e. control) data followed by the transfer of the query (i.e. case) data using the learnt model (e.g. Azimuth and scArches), would be more suitable than unsupervised integration of both reference and query data (e.g. MNN, Harmony, etc), though the performance of the latter group will depend on whether query-specific genes are included in the set of informative features used for integration.

To quantitatively assess the impact of different embedding schemes on the performance of miloDE, we analysed transcriptional changes occurring in chimeric mouse embryos, in which tdTomato+ mouse embryonic stems cells containing a Tal1 knock out were injected into wild type blastocysts (Pijuan-Sala et al. 2019). We used an atlas of Wild Type (WT; i.e. non-chimeric) gastrulating mouse embryos as a "control" (henceforth referred to as WT), and the wild type cells from the chimeric embryos were used as the "case" (henceforth referred to as ChimeraWT). Thus, we tested how the presence of Tal1- chimera cells affects ChimeraWT cells (when compared to 'true' WT cells). We computed several embeddings using both supervised (Azimuth (Hao et al. 2021), scArches (Lotfollahi et al. 2022), reference-projected MNN) and unsupervised (standard MNN (Haghverdi et al. 2018)) methods (Supplementary Note 1, Fig.1A), performing neighbourhood assignments using each approach. For the unsupervised approach, we used different highly variable gene (HVG) selections, including and excluding chimera cells, which in turn affects whether HVG selections would have genes specifically expressed in chimera embryos. To identify embedding methods that minimise the case-specific variance (in this case the chimera-specific variance), we selected 3 genes that are broadly upregulated (i.e. across the majority of cell types) in the ChimeraWT cells (Supplementary Note 1, Fig.1B) and used the standard deviation in expression across cells in a neighbourhood to assess the homogeneity of each neighbourhood. We hypothesised that in embeddings where the contribution of chimera-specific variance to the latent embedding model (and accordingly the neighbourhood assignment) is minimised, expression distributions for these genes across chimera cells from the same neighbourhoods would be randomly sampled and contain both high and low expression values. On the other hand, if chimera-specific variance contributes to the latent embedding, it is more likely that these genes contribute to the neighbourhood assignment, thus resulting in neighbourhoods that, while otherwise similar, differ in the average expression of chimera-specific genes. As expected, since the genes are chimera-specific, we observe no difference in the standard deviation of their expression in the WT cells. However, for the ChimeraWT cells, we see that the standard deviation is considerably higher for the supervised learning schemes, thus suggesting that these genes contribute less to the embeddings and are thus likely to be 'randomly' distributed across and within the assigned neighbourhoods (Supplementary Note 1, Fig.1C). Additionally, when we calculated DE for these genes between case and control samples, we also observed a systematically larger (absolute) logFC (Supplementary Note 1, Fig.1D) for the supervised embeddings. Importantly, when we compare logFC distribution across neighbourhoods to median logFC across DE tests for all abundant cell types (number of cells > 100, dashed lines in Supplementary Note 1, Fig.1D), we observe that median logFC for unsupervised embeddings is systematically lower, whereas the median for supervised embeddings mostly coincide. Overall, we suggest that supervised, reference-based embeddings will yield more sensitive DE detection and are more suitable for miloDE implementation.

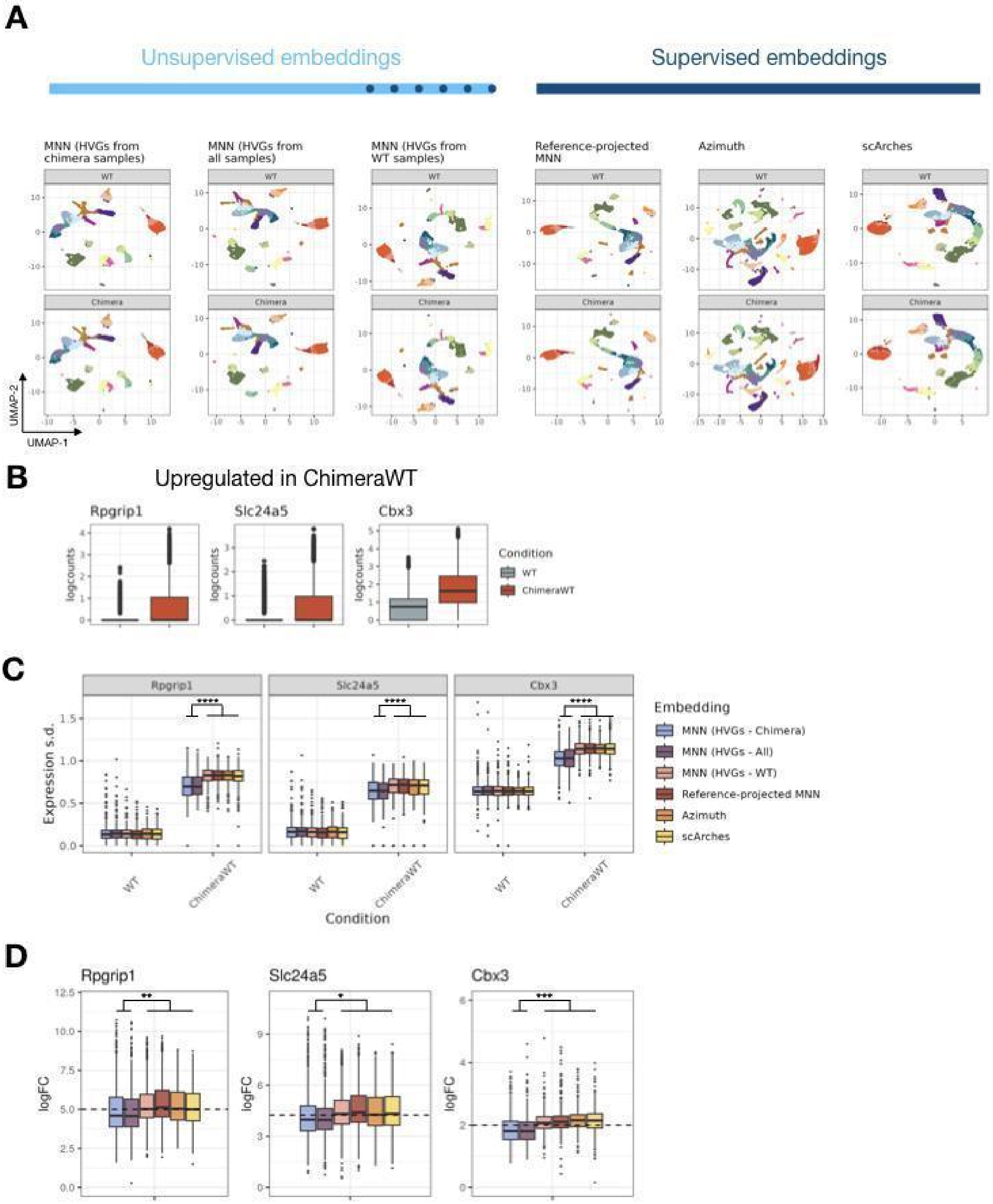


**Supplementary Note 1, Figure 1. Reference-based supervised embedding schemes yield higher sensitivity for DE identification.**

A. UMAP plots, representing different embeddings (in columns) for WT (top panels) and ChimeraWT (bottom panels) cells. Colours correspond to cell types and serve only to facilitate visualisation. We split embeddings in unsupervised (light blue) and supervised (dark blue). Note, that even though MNN embeddings are labelled as unsupervised, in case where HVG selection does not use chimera cells, chimera-specific genes are less likely to be included and therefore this embedding represents a semi-supervised approach which we indicate with dashed dark blue line.

B. Boxplots of selected chimera-specific genes, representing the expression across cells. Each facet corresponds to one gene, colours correspond to the condition.
C. Boxplots representing within neighbourhood distribution of standard deviations within the neighbourhoods for the selected genes (y-axis). Colours correspond to different embedding schemes, and facets correspond to the condition. Asterisks represent significant differences (**** < 0.00005, , Welch's T-test).
D. Boxplots representing the distribution of logFC for the selected genes. Colours correspond to different embedding schemes, and facets correspond to the condition. Asterisks represent significant differences (* < 0.05; ** < 0.005; *** < 0.0005, Welch's T-test). Dashed lines correspond to median logFC across tests within all abundant cell types (total number of cells > 100).

**Supplementary Note 2. Importance of a sufficient number of cells for sensitive DE detection.**

Since the power to detect DE using edgeR is highly dependent on the number of tested cells (Wang et al. 2019), it is important to ensure that the neighbourhood assignment method results in neighbourhood sizes (i.e. number of cells per neighbourhood) that are large enough to enable sensitive and specific DE detection. To this end, we performed simulations using `splatter` (Zappia, Phipson, and Oshlack 2017) where we varied the total number of replicates as well as the imbalance between control and case replicates, and the expected effect size (logFC) (Supplementary Note 2, Figure 1). As expected, sensitivity is highly dependent on the number of cells and replicates tested, while specificity plateaus at around 500 cells (combined across all conditions and replicates). Importantly, sensitivity to detect DE genes is, on average, below 0.75 if the number of tested cells is below 500, although this depends on the desired minimal effect size detectable (Supplementary Note 2, Figure 1C). In the original Milo approach (Dann et al. 2021), the kNN graph is used to represent the relationship between cells, and the parameter *k* (i.e. number of nearest neighbours) controls the average neighbourhood size. When tested on the mouse chimera dataset (same dataset used for the embedding analysis, Supplementary Note 1), we observed a linear relationship between *k* and average neighbourhood size: to reach an average neighbourhood size above 500, *k* needed to be above 300 (Supp. Fig. 1). More generally, while the precise relationship between *k* and neighbourhood size distribution will depend on the dataset in question, the linear dependence will force the optimal range for *k*, on average, to be on the order of hundreds. However, as *k* increases, the homogeneity of the neighbourhoods will inevitably decline, potentially leading to rare cell types (defined as those present in considerably fewer quantities than the ideal neighbourhood size) being absorbed into transcriptionally related, but more abundant cell types. To address this problem, we propose using a 2nd-order kNN graph, where we first compute the standard kNN graph (henceforth referred to as the 1st-order kNN graph), followed by assigning edges between any two cells that have at least one common neighbouring cell. To assign neighbourhoods, for each selected index cell (see Methods), we assign all cells connected with it to a single neighbourhood. As *k* increases, the average neighbourhood size increases considerably faster than it does for the 1st-order KNN graph, with a neighbourhood size in the low hundreds being achieved when *k* is between 20 and 30 (Supp. Fig. 1, right panel). Most importantly, we suggest that with 2nd-order optimization, we achieve higher neighbourhood homogeneity while controlling for average neighbourhood size. In other words, for abundant cell types that contain many transcriptionally similar cells, the method will result in sufficient neighbourhood sizes. On the other hand, for rare cell types, neighbours of neighbours will mostly lie within the same cell type, and thus the neighbourhoods will be smaller, and thus considerably more homogeneous compared to the 1st-order kNN graph.

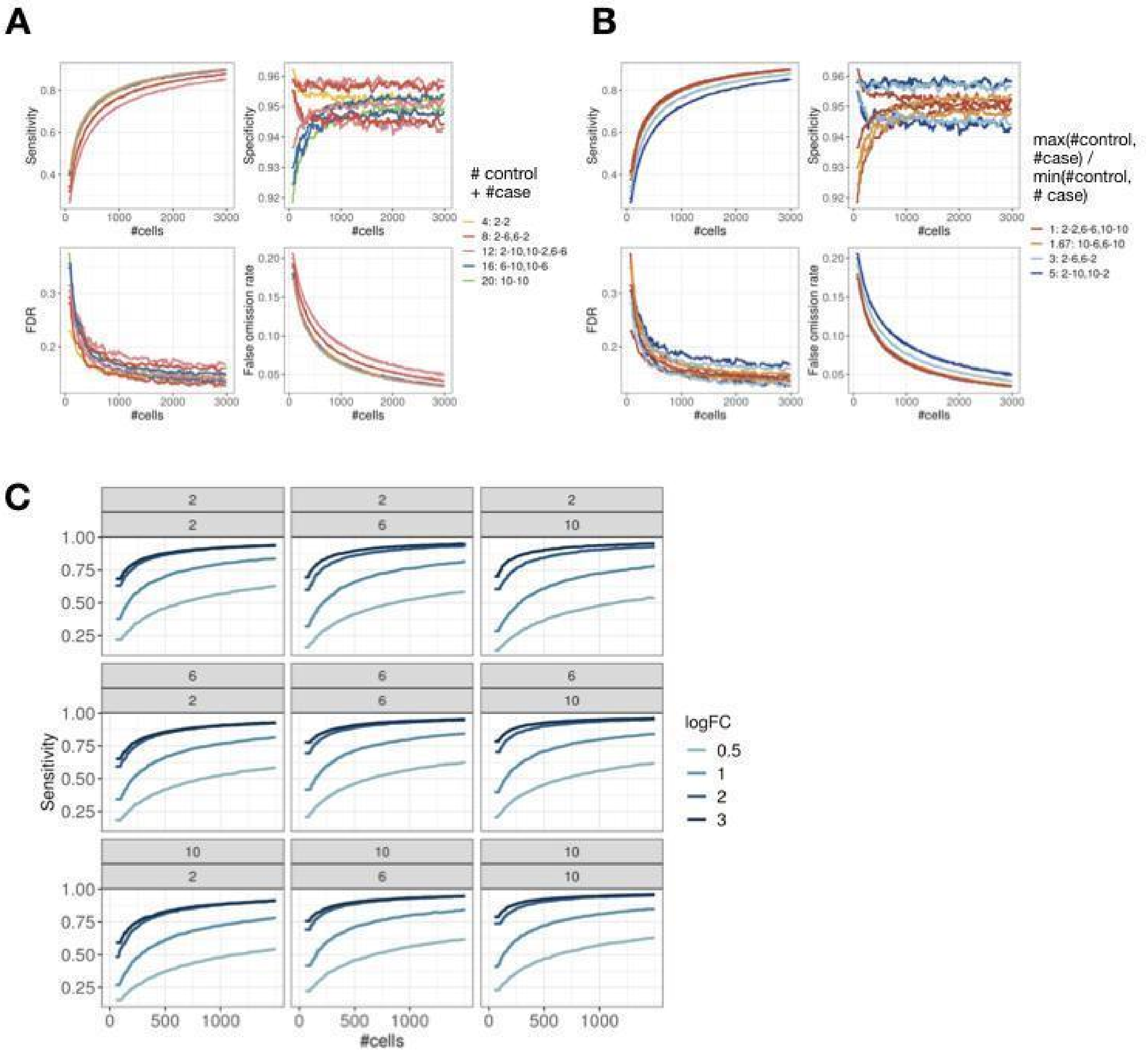


**Supplementary Note 2, Figure 1. Sensitivity in DE detection scales together with the number of replicates and number of tested cells.**

A. Trends representing the relationship between the number of cells (x-axis) and sensitivity (y-axis, top left), specificity (y-axis, top right), FDR (y-axis, bottom left) and False Omission Rate (y-axis, bottom right). Each line corresponds to one simulation; simulations vary by the total number of replicates (in colour). In the legend labels, we expand which sample composition (control and case) correspond to the total number of samples.
B. Same as A, but colours correspond to the imbalance (ratio) between the number of replicates for case and control (e.g. 1 corresponds to the same number of replicates between case and control data).
C. Trends representing the dependence of sensitivity (y-axis) on the number of cells (x-axis) and effect size (in colour). Each facet corresponds to one simulation.

**Supplementary Note 3. A priori identification of 'unperturbed' neighbourhoods decreases the number of tested neighbourhoods and reduces the burden of multiple testing correction.**

To decrease the burden of multiple testing correction, a standard practice in DE analysis is to discard genes that are lowly expressed in both case and control samples. We suggest that a similar procedure can be applied to exclude neighbourhoods from testing. To this end, we adapt the classifier algorithm Augur whose original purpose is the ranking of the cell types by the degree of their perturbation in case-control studies ((Skinnider et al. 2021), Methods). In brief, Augur builds Random Forest based classifiers to distinguish case and control cells and returns the AUC of the classifiers. We implement Augur to return the AUC for each neighbourhood, and the user can select the AUC threshold to decide which neighbourhoods will be further supplied for DE testing (recommended default is 0.5).

To test how AUC distribution depends on whether DE is present between two tested groups, we used the R package `splatter` that simulates scRNA-seq counts with the desired properties (Zappia, Phipson, and Oshlack 2017). Specifically, we generated several datasets with different number of genes that are DE (including 0 as a control), different effect size and introduced batch effect (Methods). Our simulations confirm that in comparisons where DE exists, AUCs are consistently higher than 0.5, whereas, in the absence of the DE, AUCs are driven by the existence of batch effects between case and control samples (with AUCs centring around 0.5 in datasets with no or balanced batch effect) (**Supplementary Note 3, Fig. 1**). Therefore, we suggest that while it is beneficial to discard uninteresting neighbourhoods to minimise the computing time and the burden of multiple testing correction, the practical benefit of this step in the datasets with complex batch effects is negligible, and we leave this optional step to the decision of the user. In the implementation of this step in the package, we return AUC calculated for each neighbourhood, and subsequently the user can select their own AUC threshold.

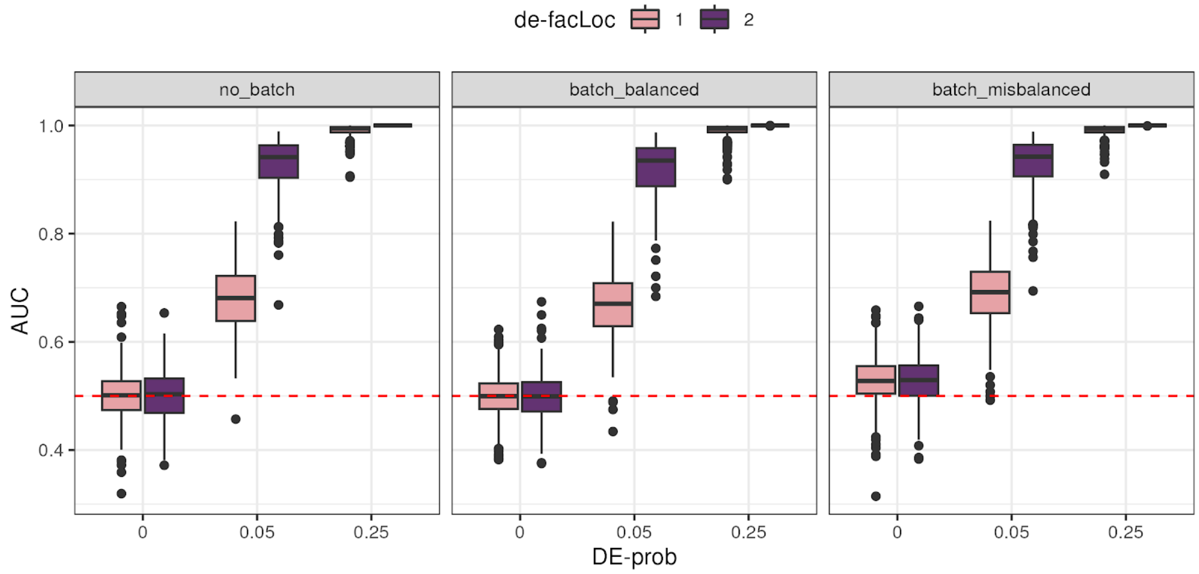


**Supplementary Note 3, Figure 1. Random Forest based classifiers successfully identify neighbourhoods with DE as 'important'.**

Boxplot representing the relationship between AUC (y-axis) of Random Forest classifiers and fraction of genes with DE (x-axis), effect size (in colour) and the existence of the batch effect (in facets). Red dashed line corresponds to AUC = 0.5 which we use to define neighbourhoods as relevant or irrelevant for DE testing.

**References.**

Chen, Huidong, Caleb Lareau, Tommaso Andreani, Michael E. Vinyard, Sara P. Garcia, Kendell Clement, Miguel A. Andrade-Navarro, Jason D. Buenrostro, and Luca Pinello. 2019. "Assessment of Computational Methods for the Analysis of Single-Cell ATAC-Seq Data." *Genome Biology* 20 (1): 241.

Dann, Emma, Neil C. Henderson, Sarah A. Teichmann, Michael D. Morgan, and John C. Marioni. 2021. "Differential Abundance Testing on Single-Cell Data Using K-Nearest Neighbor Graphs." *Nature Biotechnology*, September. https://doi.org/10.1038/s41587-021-01033-z.

Haghverdi, Laleh, Aaron T. L. Lun, Michael D. Morgan, and John C. Marioni. 2018. "Batch Effects in Single-Cell RNA-Sequencing Data Are Corrected by Matching Mutual Nearest Neighbors." *Nature Biotechnology* 36 (5): 421–27.

Hao, Yuhan, Stephanie Hao, Erica Andersen-Nissen, William M. Mauck 3rd, Shiwei Zheng, Andrew Butler, Maddie J. Lee, et al. 2021. "Integrated Analysis of Multimodal Single-Cell Data." *Cell* 184 (13): 3573–87.e29.

Lotfollahi, Mohammad, Mohsen Naghipourfar, Malte D. Luecken, Matin Khajavi, Maren

Büttner, Marco Wagenstetter, Žiga Avsec, et al. 2022. “Mapping Single-Cell Data to Reference Atlases by Transfer Learning.” *Nature Biotechnology* 40 (1): 121–30.

Luecken, Malte D., M. Büttner, K. Chaichoompu, A. Danese, M. Interlandi, M. F. Mueller, D. C. Strobl, et al. 2022. “Benchmarking Atlas-Level Data Integration in Single-Cell Genomics.” *Nature Methods* 19 (1): 41–50.

Pijuan-Sala, Blanca, Jonathan A. Griffiths, Carolina Guibentif, Tom W. Hiscock, Wajid Jawaid, Fernando J. Calero-Nieto, Carla Mulas, et al. 2019. “A Single-Cell Molecular Map of Mouse Gastrulation and Early Organogenesis.” *Nature* 566 (7745): 490–95.

Skinnider, Michael A., Jordan W. Squair, Claudia Kathe, Mark A. Anderson, Matthieu Gautier, Kaya J. E. Matson, Marco Milano, et al. 2021. “Cell Type Prioritization in Single-Cell Data.” *Nature Biotechnology* 39 (1): 30–34.

Wang, Tianyu, Boyang Li, Craig E. Nelson, and Sheida Nabavi. 2019. “Comparative Analysis of Differential Gene Expression Analysis Tools for Single-Cell RNA Sequencing Data.” *BMC Bioinformatics* 20 (1): 40.

Zappia, Luke, Belinda Phipson, and Alicia Oshlack. 2017. “Splatter: Simulation of Single-Cell RNA Sequencing Data.” *Genome Biology* 18 (1): 174.
